# Exploring lipid nanoparticle design spaces using self-regulating microfluidic machines and multiplexed *in vivo* biodistribution

**DOI:** 10.64898/2026.06.03.729611

**Authors:** Josef Kehrein, Elena Reus, Caroline T. Holick, Sebastian Mummel, Ilayda Ates, Markus Hafke, Jonas Käsbach, Christine Weber, Tessa Lühmann, Stephanie Schubert, Jørgen Magnus, Florian A. Mann, Ulrich S. Schubert, Lorenz Meinel

**Author notes:** Both authors contributed equally to this work (Josef Kehrein, Elena Reus).

## Abstract

Delivering therapeutic mRNA relies on lipid nanoparticles (LNPs). Finding optimal process parameters for new lipid combinations in LNP formulations remains a challenge. In our work, we used an automated, self-regulating microfluidic platform that actively changes process parameters to tune LNP formulations for preset, desired quality standards. We tested new LNPs by swapping poly(ethylene glycol) (PEG) lipopolymers for alternatives based on poly(2-methyl-2-oxazoline) (PMeOx) and poly(2-ethyl-2-oxazoline) (PEtOx). For each lipopolymer variant, the platform independently identified optimal production conditions in four or fewer iterative cycles, yielding particles of the preset size and high mRNA encapsulation. Small-angle X-ray scattering revealed that smaller LNPs modified with PEtOx had more structural surface variety and higher mRNA loading efficiency. When multiplexing these formulations in mice, the PEtOx-containing LNPs accumulated more in bone marrow compared to those with PEG, indicating trends that the chemistry of the lipopolymer affects the biodistribution of the resulting LNPs. By combining automated formulation and *in vivo* multiplexed testing, our approach provides a practical way to rapidly plan, formulate, and evaluate large pharmaceutical design spaces, to select excipients and process parameters yielding optimal biological performance of LNPs.

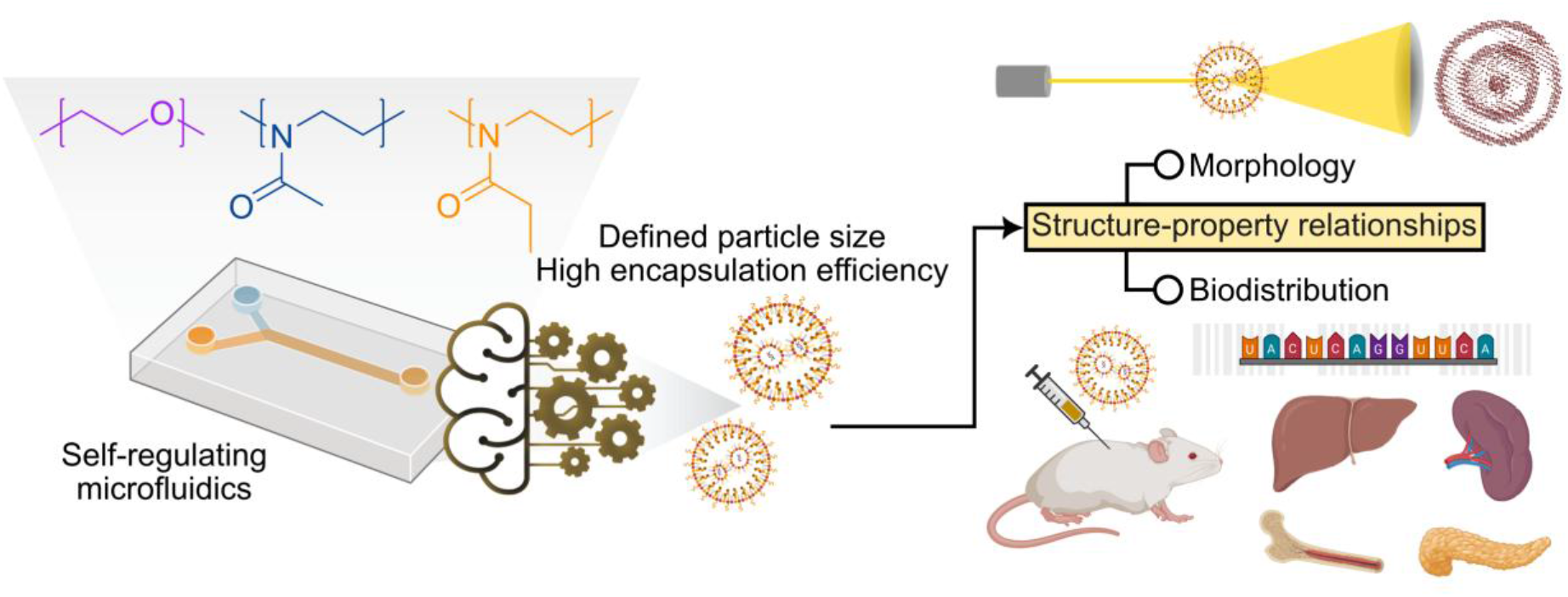

## 1 Introduction

Lipid nanoparticles (LNPs) have evolved into gene delivery systems across various therapeutic fields [1–3]. Current research on these vehicles is guided by the formulation parameters and lipid compositions of the few marketed LNPs, specifically the COVID-19 mRNA vaccines Elasomeran/Spikevax® (Moderna) and Tozinameran/Comirnaty® (Pfizer/BioNTech) [4, 5]. Many studies reported on the effects of exchanging phospholipids [6], cholesterol [7], ionizable aminolipids [8], or the stealth lipopolymer containing poly(ethylene glycol) (PEG) [9, 10]. These works are essential to overcome central pharmacokinetic obstacles, such as emanating from hepatocellular accumulation [11]. However, such endeavors entail time-and resource-intensive experimental screening, as even subtle changes in process settings or excipients can result in pronounced shifts in critical quality attributes (CQAs) and biological activity [12–14]. In principle, whenever new lipid components are introduced, e.g., intended for targeting tissues beyond the liver, or another nucleic acid cargo is loaded, the process parameters driving LNP formation will require adaptation. Regarding microfluidics, which is a common production technique for LNPs, different total flow rates (TFR), flow rate ratios (FRR) between ethanolic lipid and aqueous mRNA phases, the total lipid/mRNA concentrations prior to mixing and ratios between ionizable lipids and nucleotides (N/P) are effective means to tune formulation outcomes [15] and have been used by self-regulating microfluidic devices for user-independent optimization of lipid nanoparticle formulations (**Figure 1A**) [16]. This self-regulating microfluidic device produced 95% of particles with the target size, requiring a maximum of three formulation iterations to reach it.

**Figure 1:**
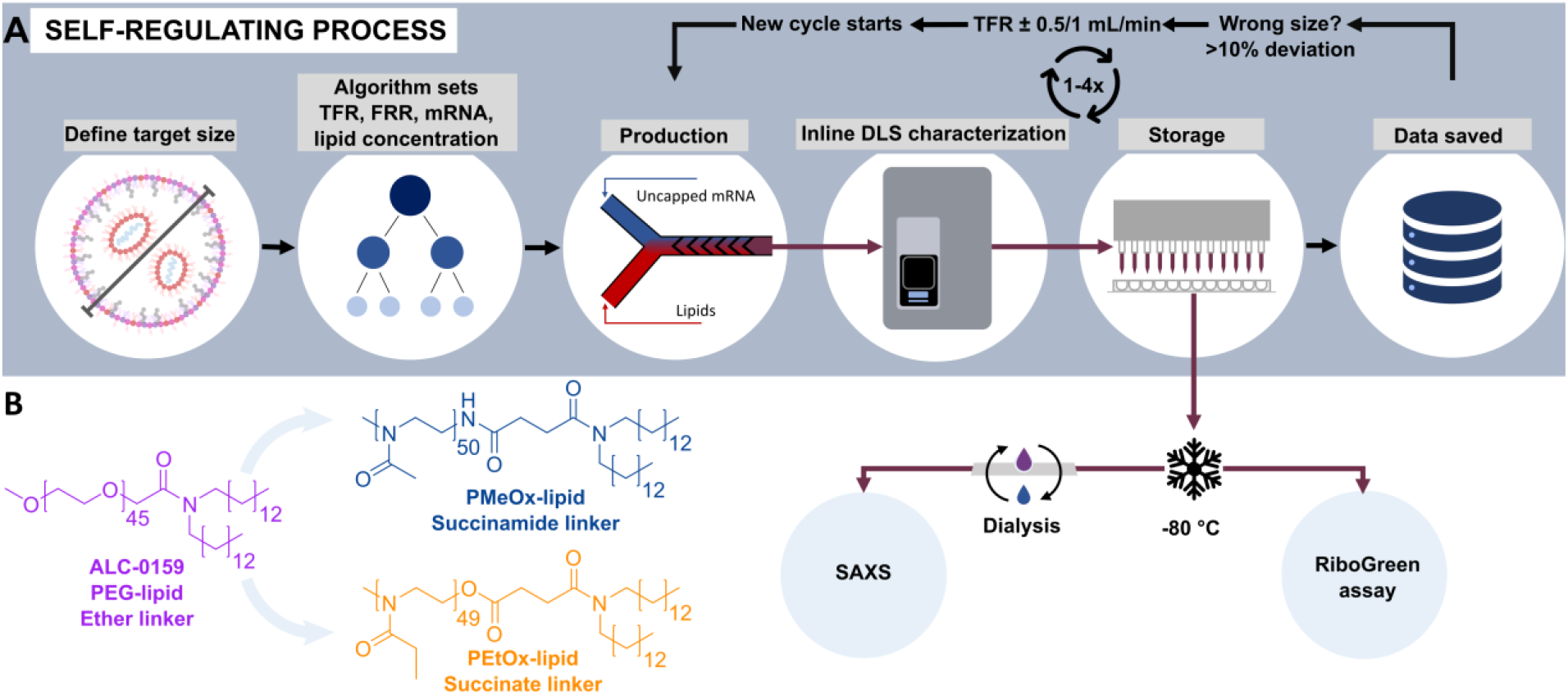
Self-regulating microfluidic device for LNP production. (A) In our previous work, we developed a device that first determined optimal formulation settings for a given target particle size, using a random forest prediction model. Subsequently, LNPs were produced and automatically characterized in-line by dynamic light scattering (DLS) [16]. If the experimentally determined size was not within a given tolerance range, the setup adjusted the TFR and formulated a new batch of particles. This loop was repeated a maximum of four times to successively reach the desirable size through optimization of process parameters. (B) Schematic representation of the PEG- and POx-lipids used in this study.

We are now expanding to alternative excipients than PEG, i.e., poly(2-methyl-2-oxazoline) (PMeOx) and poly(2-ethyl-2-oxazoline) (PEtOx), and integrate the self-regulating formulation setup with multiplex-biological assessment, to streamline high-throughput characterization of large design spaces and their biological potential. POx-based materials have shown excellent biocompatibility, as such [17, 18] or when integrated in bioconjugates [19–25]. POx-lipids have already been used for LNP production and are regarded as well suitable alternatives to PEG-lipids [26–33], which are increasingly viewed critically [34, 35]. However, a switch from PEG-to POx-based materials may substantially impact formulation outcomes [33]. For example, using a molar mass-equivalent PEtOx chain resulted in approximately double the particle size of an analogous PEG-based formulation [29]. The issue poses an interesting task for the self-regulating microfluidic formulation device, which was challenged to autonomously (without human intervention) meet preset lipid nanoparticle characteristics and rapidly map pharmaceutical design spaces in this study. Selected formulations were characterized for mRNA encapsulation efficacy (EE) and for morphology by small-angle X-ray scattering (SAXS) [36]. The broad formulation spaces provided by the self-learning machine were continued with pharmacokinetic multiplexing studies, using *in vivo* barcoding study in mice.

## 2 Materials and methods

### 2.1 Polymer synthesis

The synthesis of PEtOx-lipid was performed as previously described [33, 37]. Detailed procedures including the synthesis of PMeOx-lipid and characterization data can be found in the Supplementary Information (**Scheme S1-2, Table S1-2, Figure S1-4**).

### 2.2 Lipid nanoparticle formulation

#### 2.2.1 Materials

CatPure™ eGFP uncapped mRNA from CATUG Life Technology Co., ALC-0159 (α-(2-(ditetradecylamino)-2-oxoethyl)-ω-methoxy-poly(oxy-1,2-ethanediyl)), ALC-0315 ([(4-hydroxybutyl)azanediyl]di(hexane-6,1-diyl)bis(2-hexyldecanoate)), and cholesterol were kindly provided by Evonik Industries AG (Essen). 1,2-Distearoyl-*sn*-glycero-3-phosphocholine (DSPC) was bought from Avanti Polar Lipids, Inc.. For determination of EE, Tween 20, Tris-EDTA (TE), and the Quant-it™ RiboGreen RNA™ Assay-Kit were purchased from Thermo Fisher Scientific.

#### 2.2.2 Formulation parameters

LNPs were produced based on the Comirnaty lipid composition (46.3:42.7:9.4:1.6 molar ratio of ALC-0315, cholesterol, DSPC, and ALC-0159), whereby ALC-0159 was exchanged at 1.6 mol% with PMeOx-lipid or PEtOx-lipids. Initial settings for lipid and mRNA concentrations, as well as FRR and TFR values were constrained by our microfluidic setup (**Table S3**). Uncapped eGFP mRNA was utilized as cargo material. 20 different randomly selected target sizes below 120 nm were defined for both polymer types, allowing the applied random forest prediction model (*vide infra*) to select the best-performing combination of settings. Subsequent runs, in which our self-regulating mechanism was triggered, were altered by changing TFR values either by ± 0.5 or ± 1 mL/min (depending on the measured value). After production and DLS characterization, particles were further diluted 1:3 with a phosphate buffered saline (PBS)-glycerol mixture and frozen at 80 °C, resulting in a final glycerol concentration of 30% until further usage for investigations *via* RiboGreen assay and SAXS measurements.

Particles selected for SAXS measurements from each run were dialyzed to remove the cryoprotectant, ethanol, acetate buffer, and any remaining impurities potentially affecting measurements. Dialysis was performed using a 0.5 mL dialysis tube with a 10 kDa molar mass cutoff (Slide-A-Lyzer®, Thermo Fisher Scientific). The dialysis tube was filled with 0.5 mL of LNP suspension and dialyzed against 13 mL of PBS under continuous stirring for 3 h at room temperature, and then overnight at 5 °C. During the first 3 hours, the buffer was changed every hour. The SAXS measurement took place one day after dialysis.

#### 2.2.3 Hardware setup

LNPs were produced with a Fluidic 187 micromixer (Microfluidic Chipshop), as described previously (see also **Figure 5** in [16] for a more detailed overview). Briefly, a staggered herringbone micromixer served as mixing device, whereas the fluid flow was regulated by FLOW EZ pressure and L/L+ flow units. Lipid and nucleotide concentrations were varied using 10-port M-switches. Buffer solutions were used for flushing and stabilization. The LNP stream was diluted 1:1 with PBS by a T-mixer, with subsequent particle size analysis by a ZetaSizer Ultra-Red machine (see below). LNPs were dispensed by a WELLJET reagent dispenser into 96-well plates for further experiments on EE and easier storage.

#### 2.2.4 Software setup

The same software setup as in our previous study was applied [16]. In short, we controlled our microfluidic device using the OxyGEN 2.1.0.0 software combined with in-house python scripts, automatically regulating pressure and flow controllers and all valves within the system. This allowed controlling all individual steps of LNP formulation in sequence or parallel (washing, production, data generation, and particle collection). Data on particle size were collected by DLS measurements in-line, using the ZS Xplorer software (version 1.32) of the ZetaSizer Ultra-Red machine. The self-regulation mechanism was implemented by first setting a target LNP size with a tolerance threshold of 10% within our python script. Based on the desired value, our previously generated size prediction model (*vide infra*) was utilized to find optimized formulation parameters. After assessing particles *via* DLS, the most important feature variable of our model, the TFR, was adjusted if the tolerated interval was not reached. In this case, a second run with an updated TFR setting was performed and new particles were characterized again (± 0.5 or 1 mL/min, for when the tolerated range was exceeded by 10 to 20% or >20%, respectively). This self-regulating mechanism was run automatically within our microfluidic setup for a total of 2 to 3 formulation repeats (i.e., up to 3 total production cycles for the PEG-lipid and PMeOx-lipid, and four total runs for the PEtOx-lipid), successively adjusting the TFR to meet the desired target size.

### 2.3 Lipid nanoparticle characterization

#### 2.3.1 Dynamic light scattering

A ZetaSizer Ultra-Red (Malvern Panalytical) was used for determining LNP particle sizes (Z-average hydrodynamic diameters, D_H_) and polydispersity index (PDI) values. For in-process DLS measurements, a ZEN2112 cell at 25 °C with back scatter angle detection, automatic attenuation, measurement at a fixed position, no optical filter, a refractive index of 1.33, and a viscosity of 1.2824 mPa s (equaling 10% ethanol) was used, applying the Smoluchowski approximation. 30 runs were performed, each at a duration of 1.68 s, with an initial equilibration time of 10 s and no pause in-between runs. Measurements were made in technical triplicate. As previously described [16], the LNP sizes were subsequently re-calculated, by relating size and viscosity for 10% ethanol to the real ethanol content based on the Stokes equation. For particle size determination after dilution, a DTS0012 cell with a set viscosity of 0.8872 mPa s was used, measuring at optimal position. All other settings were the same as mentioned for the in-process measurements.

#### 2.3.2 Encapsulation efficiency

Total and encapsulated mRNA were quantified as described previously [16], using the RiboGreen™ RNA Assay. A 6-point calibration curve (0.02 – 1 μg/mL mRNA) was used for fluorescence-based concentration measurements. Encapsulated mRNA was calculated by subtracting unencapsulated (measured without detergent) from total mRNA (measured after LNP lysis with Tween 20). EE was expressed as encapsulated/total mRNA (%). Samples were diluted in TE buffer, prepared in black 96-well plates with blanks included, and analyzed in triplicates, with the mean value reported for results. After incubation with Tween 20 (37 °C, 10 min) and RiboGreen (5 min, room temperature, dark), fluorescence was measured at 485/530 nm using a microplate reader (BioTek Synergy MX).

#### 2.3.3 Size prediction model

The regression model used for prediction of particle sizes within the self-regulating mechanism was a random forest previously built via sci-kit learn [38], based on LNPs with ALC-0159 loaded with CatPure™ eGFP uncapped mRNA [16]. The independent feature variables used were the TFR, the N/P ratio, the nucleotide concentration, the lipid concentration, and the FRR. To find initial parameter settings for a given size, the device ran the implemented prediction model for all allowed combinations of TFR, FRR, and mRNA/lipid concentrations (according to **Table S3**). The model input N/P ratio was derived from the mRNA/lipid concentration and the FRR. Subsequently, settings of the prediction nearest to the target size were set for the initial production run.

#### 2.3.4 Small angle X-ray scattering

To study the morphology of LNPs with alternative lipopolymers, SAXS measurements were performed at the PETRA III synchrotron (P12 beamline) at DESY (Hamburg, Germany). Therefore, we formulated a single batch of unloaded and five different batches of loaded PEtOxylated LNPs with different target sizes using our microfluidic setup (**Table S4**).

SAXS data (*I*(*s*)) *vs. s*, where the momentum transfer is 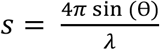, 2θ is the scattering angle, and *λ* = 0.124 *nm* is the X-ray wavelength) were collected with the undulator (gap 20.95 mm) and monochromator adjusted for 10 keV. The beamline was calibrated for angular axis using silver behenate powder. Prior to measurements, the beamline was tested [39, 40]: measurements of empty capillary, Milli-Q® water, bovine serum albumin (BSA), and HEPES buffer (4-(2-hydroxyethyl)-1-piperazineethanesulfonic acid) were performed in batch mode (sample loading with a sampler changer robot). Further specifications are listed in **Table S5**. Samples were handled with an autosampler and measured in batch mode [39, 40]. Here, the samples were loaded to an in-vacuum flow-through quartz capillary for exposure with X-rays. To limit radiation damage, samples were continuously flown when exposed to the X-ray beam. The storage tray was temperature-controlled and set to 20 °C. For each sample at each concentration, 40 successive images were collected. The images were compared and only similar images with no radiation damage were averaged for further analysis. To isolate the scattering contribution of the samples, scattering of Milli-Q® water was used for background subtraction. 40 background images were collected before and after each sample measurement. The collected scattering data underwent standard automated radial averaging [39, 40], and the data were processed and analyzed using the ATSAS program suite [41]. The overall particle parameters (radius of gyration, *R*_*g*_, and maximum size, *D*_*max*_) were assessed by PRIMUS [42] with Guinier approximation [43] according to **Equation (1)**. The program GNOM was utilized to compute pair distribution functions *p(r)*, according to **Equation (2)** [44]. The ordering in the samples was evaluated by the program PEAK [41, 42], according to **Equations (3)** and **(4)**, where *s*_*max*_ is the momentum transfer of the peak, *λ* represents the wavelength of X-rays, *β*_*s*_ the full width at half-maximum intensity of the peak (in radians), and θ_*max*_ the scattering angle corresponding to *s*_*max*_. Additionally, shape determination was conducted by the *ab initio* bead modeling program DAMMIN [45].

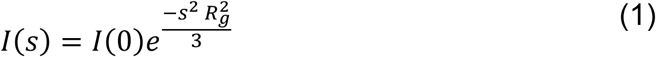

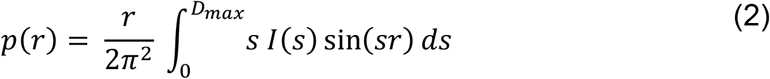

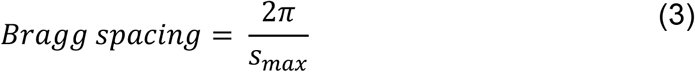

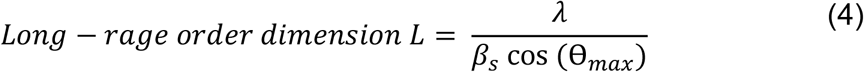

### 2.4 In vivo biodistribution analysis

#### 2.4.1 LNP formulation and preparation

LNPs for the barcoding experiment were formulated at a concentration of 30 mM lipid in a 10 mM citric buffer at pH 4, using a microfluidic mixing setup at a flow rate of 12 mL/min. Lipids were dissolved in ethanol and rapidly mixed with an aqueous phase containing nucleic acids, using the NanoAssemblr™ platform (Precision NanoSystems). The aqueous phase contained a short single-stranded DNA barcode specific to each formulation and a Firefly luciferase (FLuc) mRNA. Following formulation, the LNPs were dialyzed against PBS to remove unencapsulated materials and ethanol. The final product was sterile filtered using a 0.2 µm poly(ether sulfone) (PES) filter. Particle size and PDI value were measured by DLS and ζ-potential by electrophoretic light scattering (ELS), using a Zetasizer Nano Series ZS instrument (Malvern Panalytical). For DLS measurements, a ZEN0040 or Fisher 759200 cell at 25 °C with an equilibration time of 60 s was used, applying back scatter angle detection, automatic attenuation, measurement at a fixed position, no optical filter, a refractive index of 1.35, and a viscosity of 1.2 mPa s (equaling 5% Tris-Sucrose), based on the Smoluchowski approximation. 30 runs were performed, each at a duration of 1.68 s with no initial equilibration time and no pause in-between runs. EE was determined as described previously [16], using the RiboGreen™ RNA Assay according to the manufacturers’ instructions.

#### 2.4.2 LNP cell toxicity

Prior to animal studies, toxicity was assessed by ATP viability using a CellTiter-Glo® 2.0 Cell Viability Assay (Cat. G9243, Promega), with readout performed on a BioTek Synergy Neo2 Hybrid Multimode Reader (Agilent). HepG2 cells were seeded onto 96-well plates (10,000 / well, 200 µL / well) and incubated for 24 h prior and post-treatment with LNPs. Measurements were performed in triplicate and values normalized between the negative (100%) and the dead control (0%).

#### 2.4.3 In vivo biodistribution screening

To enable parallel evaluation of multiple formulations, a pooled *in vivo* screening approach based on DNA barcode tracking was used as previously described [46–48]. Briefly, 38 different LNP formulations, each encapsulating a unique DNA barcode oligonucleotide, were pooled at equal ratios prior to *in vivo* administration. Adult C57/BL6 mice were treated with intravenous injections of the pooled LNP library (1 mg/kg). Our study included two groups with five animals each and two control animals. At 6 (group 1) and 24 hours post-injection (group 2), tissues including liver, heart, spleen, lung, kidney, brain, bone marrow, and whole blood were harvested and genomic DNA was extracted. Barcode sequences corresponding to each LNP formulation were amplified by polymerase chain reaction (PCR) and quantified by next-generation sequencing to determine relative tissue distribution. While our pooled screening included in total 38 LNP formulations, only LNPs decorated either with the PEtOx-lipid discussed within this work or with one of the PEG benchmarks (ALC-0159 from Comirnaty and DMG-PEG2000 from Spikevax) were analyzed and reported here.

## 3 Results and discussion

As previous studies conducted by various research groups have found that POx with lipid end groups were suitable for LNP formation if the degree of polymerization (DP) was similar as that of PEG_2k_, which is used in ALC-0159, we targeted DP values around 50 for the synthesis of the POx-lipids [26, 28, 29, 32]. For PEtOx with a DP of 50, this results in a similar hydrodynamic volume as that of PEG_2k_ [49]. As the hydrodynamic volume of PMeOx is larger than that of PEtOx at constant molar mass, we kept the DP value similar to achieve a similar hydrodynamic volume of both POx. The POx-lipids differed in the linker functionality: Whereas both are based on a succinoyl spacer, the linker is attached *via* an amide moiety to PMeOx but *via* an ester moiety to PEtOx (**Figure 1B**). In consequence, synthetic routes differed: Following a previously reported four-step procedure (**Scheme S1**) [37], PEtOx-lipid was obtained through a series of post-polymerization modifications at the *ω*-acetate end group, which resulted from termination of the cationic ring-opening polymerization (CROP) of 2-ethyl-2-oxazoline with triethylammonium acetate. In contrast, the CROP of 2-methyl-2-oxazoline was terminated with sodium azide. After Staudinger reduction of the *ω*-azide end group yielding the primary amine, coupling with 4-(ditetradecylamino)-4-oxobutanoic acid gave the desired PMeOx-lipid (**Scheme S2**). The success of the various end group transformations was demonstrated by the respective end group signals in the ^1^H nuclear magnetic resonance spectra (**Figure S1A, S3A**). Their covalent attachment was confirmed by matrix assisted laser desorption / ionization time-of-flight mass spectrometry (**Figure S2, S4**). According to characterization by means of size exclusion chromatography (**Figure S1B, S3B**), molar mass distributions of the final POx-lipids were narrow and unimodal, with dispersity values Đ of 1.18 for PMeOx-lipid and Đ of 1.05 for PEtOx-lipid. Further characterization data can be found in the Supplementary Information (**Table S1-2, Figure S1-4**).

### 3.1 Formulating POxylated LNPs by a self-regulating microfluidic machine

We selected 12 and 9 different target sizes for PMeOx- and PEtOx-lipids, respectively. For PEGylated LNPs, using the Comirnaty PEG-lipid ALC-0159, our previous investigations included ten different target sizes, and these previously reported data sets are used for comparison again in this work [16]. The random forest regression model developed for the PEGylated LNPs required self-regulated repetitions to reach target sizes in 40% of runs, or, in other words, in 60% of cases preset target diameters were immediately hit with the first production cycle (**Table S6**) [16]. That algorithm, developed for PEG-lipids, was now expanded to formulate POx-lipids. In these runs, we first allowed the self-regulating machine to undergo a maximum of two adaptations of its production process. When using PMeOx-lipid, four batches with each having another preset target value required a single adaptation in production settings, and another preset size required two adaptations to reach the tolerated interval of 90 to 110% of the preset diameter (**Figure 2, Table S6**). A single target (107 nm) was not reached within the set limitation of two adaptations. For the PEtOx-lipid, process adaptations were triggered 6 out of 9 times (67%), with a single adaptation needed for three batches and two adaptations required for the target value of 65 nm. The preset size of 83 nm was only narrowly missed (10.59% deviation) within the two adaptations allowed. However, when increasing the limit of adaptations to three (user-definable), as done exemplarily here for the preset size of 91 nm, the desired target value was reached in the additional production cycle.

**Figure 2:**
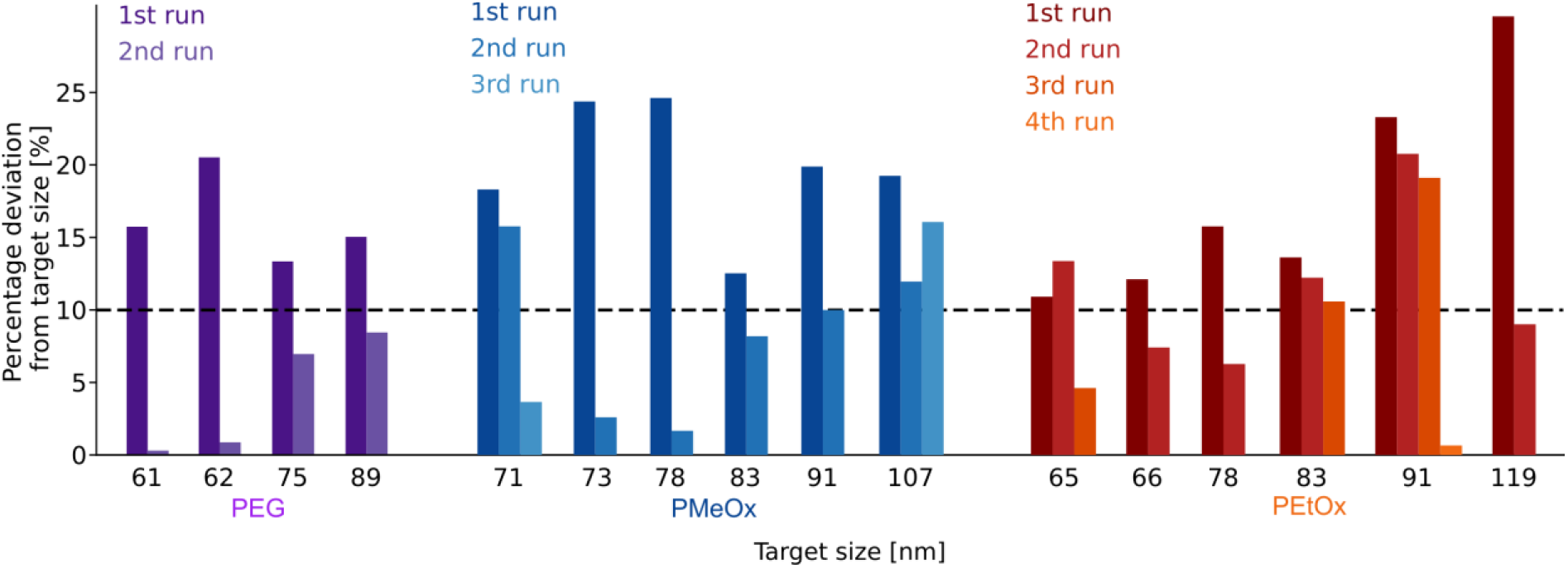
Self-regulation mechanism results. Particle size deviations from target values after every production cycle. Only runs where the initial production cycle did not reach the respective target size are shown (all runs are listed in **Table S6**). Size deviations are depicted for PEGylated (violet, adapted from [16], licensed under CC-BY), PMeOxylated (blue) and PEtOxylated LNPs (orange).

As mentioned, the self-regulating machine’s algorithm was developed with PEG-lipids [16]. Consequently, even though multiple runs directly reached the target size (**Table S6**), the machine’s first production cycles conducted with POx-lipids more often resulted in different sizes than what its algorithm (trained on PEG-lipids) would have expected, and the machine independently entered into repetitive adaptation of process parameters to reach preset target diameters. Thereby, the machine independently found its own way in the pharmaceutical design space, or “trajectory”, to reach target sizes. We visualize these trajectories, or the path the machine found to reach target sizes, by a principal component analysis (PCA; **Figure 3A, Table S6**). One area, characterized by relatively high diameters (target sizes of 87 to 105 nm) and high mRNA and lipid concentrations, was consistently identified by the algorithm in its first production cycle, independent of polymer choice (**Figure 3A**, yellow circle). In this region, deviations from larger target dimensions may be tolerated better than for smaller LNPs, as the self-regulating machine accepts batches based on percentages of the target diameters; consequently, larger LNPs allow larger tolerated limits in absolute terms than smaller particles. Our analysis also revealed that PEG-lipid and PMeOx-lipid formulations tended to yield particles larger than 110% of the target value. By contrast, PEtOxylated LNPs were usually smaller than 90% of the target value (**Table S6**). We speculated that the additional methylene group incorporated into the bulkier, more hydrophobic PEtOx side chain, compared to PMeOx, may reduce the polymer content per particle required to shield the same area of the LNP surface. That would result in different surface-to-volume ratios and systematically smaller particles at otherwise identical process settings [9, 50]. We visualized both the influence of the choice of polymer and the target size on the size deviation per cycle (**Figure 3B**). Here, the convergence of PEGylated and PMeOxylated particles to the desired interval starting from >110% and PEtOxylated particles starting from <90% becomes evident, with most particles requiring only a single production run to reach target sizes when these target sizes ranged between 95 to 120 nm.

**Figure 3:**
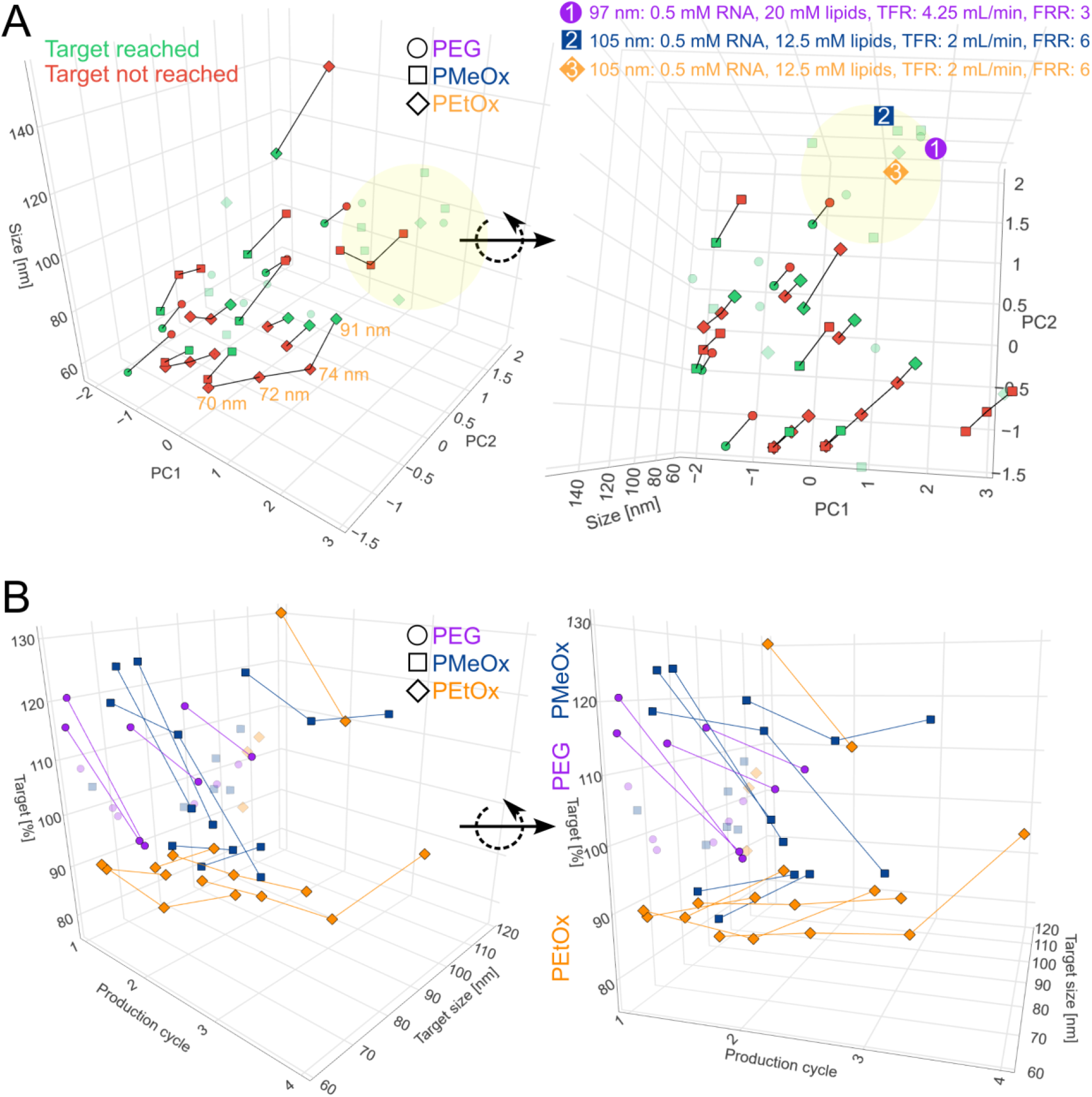
Self-regulation mechanism trajectories. (A) Experimentally determined sizes (Z axis) plotted against the experimental settings that were summarized in the form of two principal components (PC1 and PC2) after PCA on all parameters. Points within the tolerance threshold (90 to 110% of the target size) are colored green, those above or below red. For instances where the mechanism was triggered, consecutive measurements are connected by black lines, thereby illustrating the “trajectory” through the experimental design space. Instances that did not trigger the mechanism are shown as green transparent points. For an exemplary PEtOx-lipid trajectory with the target size of 91 nm, diameters obtained at each production cycle are shown in orange. Additionally, a design space area encompassing many “first-run positives” is highlighted in yellow. (B) Percentage deviations of measurements plotted against production cycle and target size. Here, points are colored according to the lipopolymer variant. Instances that did not trigger the mechanism are shown as transparent points.

The self-learning machine successfully (and independently) expanded its initially PEG-lipid-trained algorithm to meet preset LNP target sizes when using PMeOx-lipid and PEtOx-lipid in LNPs. The approach demonstrated that LNPs with preset target sizes were derived from these polymers within relevant diameters (60 to 120 nm) and the particles had acceptable PDI values below 0.25 [51, 52].

### 3.2 Encapsulation efficiency and LNP structure

EE decreased in the order of PEG-lipid > PMeOx-lipid > PEtOx-lipid (**Figure 4, Table S6**). Smaller LNPs were more likely to exhibit high EE (near 80% or above), regardless of the lipopolymer used for formulation, compared to larger-sized particles. Interestingly, at very large sizes (>120 nm), PMeOxylated LNPs retained high EE values of around 80%, whereas particles formulated with the slightly more hydrophobic PEtOx-lipid had substantially lower loadings of around 40%. Overall, our observations regarding EE suggest that, in addition to the hydrophilic polymer “corona” forming on the outer surface of LNPs, the choice of lipopolymer affected mRNA packing and possibly LNP structure. To verify differences in LNP structure, we further analyzed the morphology of PEtOxylated particles using SAXS, as PEtOx resulted in the largest difference in EE compared to PEGylated LNPs (with the latter already having been characterized *via* scattering experiments in various studies [36, 53–55]).

**Figure 4:**
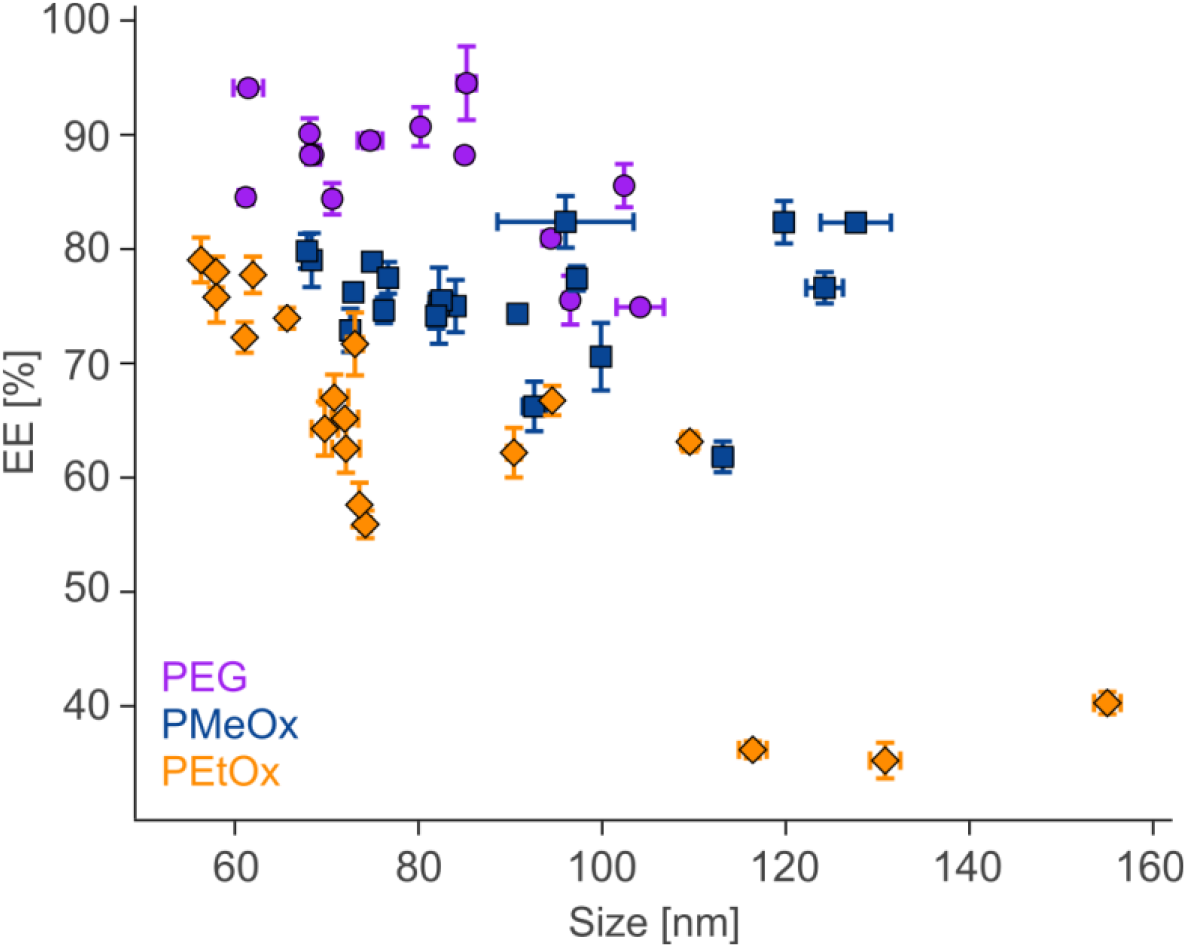
Encapsulation efficiencies. The plot shows EE values of all particles produced by our self-regulating device, depending on their sizes as determined by DLS (**Table S6**, mean ± SD, n=3).

For that, PEtOxylated LNPs of five different sizes were formulated using the automated device (samples named PEtOx-LNP1-5, **Table S4**), ranging from 56 to 73 nm in size with N/P ratios of 5 to 19. Additionally, unloaded LNPs (PEtOx-LNP6) were produced. Subsequent SAXS measurements yielded scattering curves with a rather sharp intensity increase at the smallest angles and little structure at higher angles (**Figure 5A**). All samples exhibited a shoulder around *s* = 0.4 nm^-1^, pointing to isometric shapes. For loaded particles, SAXS-derived d_max_ values (**Table S7**) correlated with DLS measurements obtained in-line by the self-learning microfluidic machine (**Figure 5B**). DLS outcomes were systematically about ~20 nm larger compared to SAXS outcomes, reflecting that DLS-measured hydrodynamic diameters also accounted for hydrated surface layers of LNPs [56–58]. Interestingly, pair distribution functions *p(r)* for samples PEtOx-LNP1-3 resulted in negative values at higher distances (**Figure 5C**). Such negative excursions, to our knowledge, were not yet observed for PEGylated LNPs and, therefore, could result from structural changes introduced by the PEtOx-lipid when formulated into smaller-sized mRNA-loaded particles (in-line d_H_ values of 52 to 58 nm). These excursions may indicate either repulsive interactions between particles in suspension or inhomogeneous density distributions at the particle periphery. However, as values increased again at the highest distances, density changes are the more likely reason for this observation. The outer monolayer of LNPs is mainly composed of cholesterol (~63 mol% in Comirnaty LNPs, near the excipient’s solubility limit [59]), which shows a lower electron density compared to water (**Table S8**). In contrast, PEtOx-lipid and DSPC molecules (with the latter being present on the surface but also involved to small degrees in internal mRNA packing [59]) both exhibit higher densities. Therefore, the negative excursions in *p(r)* profiles could originate either from a higher abundance of DSPC molecules at the outer periphery at larger N/P ratios (10 to 19), where mRNA vesicles may be solely composed of aminolipids, or from an increased mixing of the outer cholesterol-rich phase with polymers, thereby shifting the overall electron density towards a value close to that of water. A previously reported increase in polymer-cholesterol interactions in PEtOxylated particles corroborates these mixing tendencies [60], and other studies have also employed SAXS to successfully detect surface structural changes in LNPs [10, 53, 61, 62]. Larger unloaded particles (PEtOx-LNP6, d_H_ = 70 nm) exhibited no such *p(r)* curves. Furthermore, the distribution functions of mRNA-loaded PEtOx-LNP4 and PEtOx-LNP5 samples (d_H_ of 66 and 73 nm, respectively) also showed a non-negative, much more skewed profile, in accordance with the higher PDI values determined by our formulation setup (**Table S4**). For monodisperse systems, such profiles, in principle, would point to elongated shapes.

**Figure 5:**
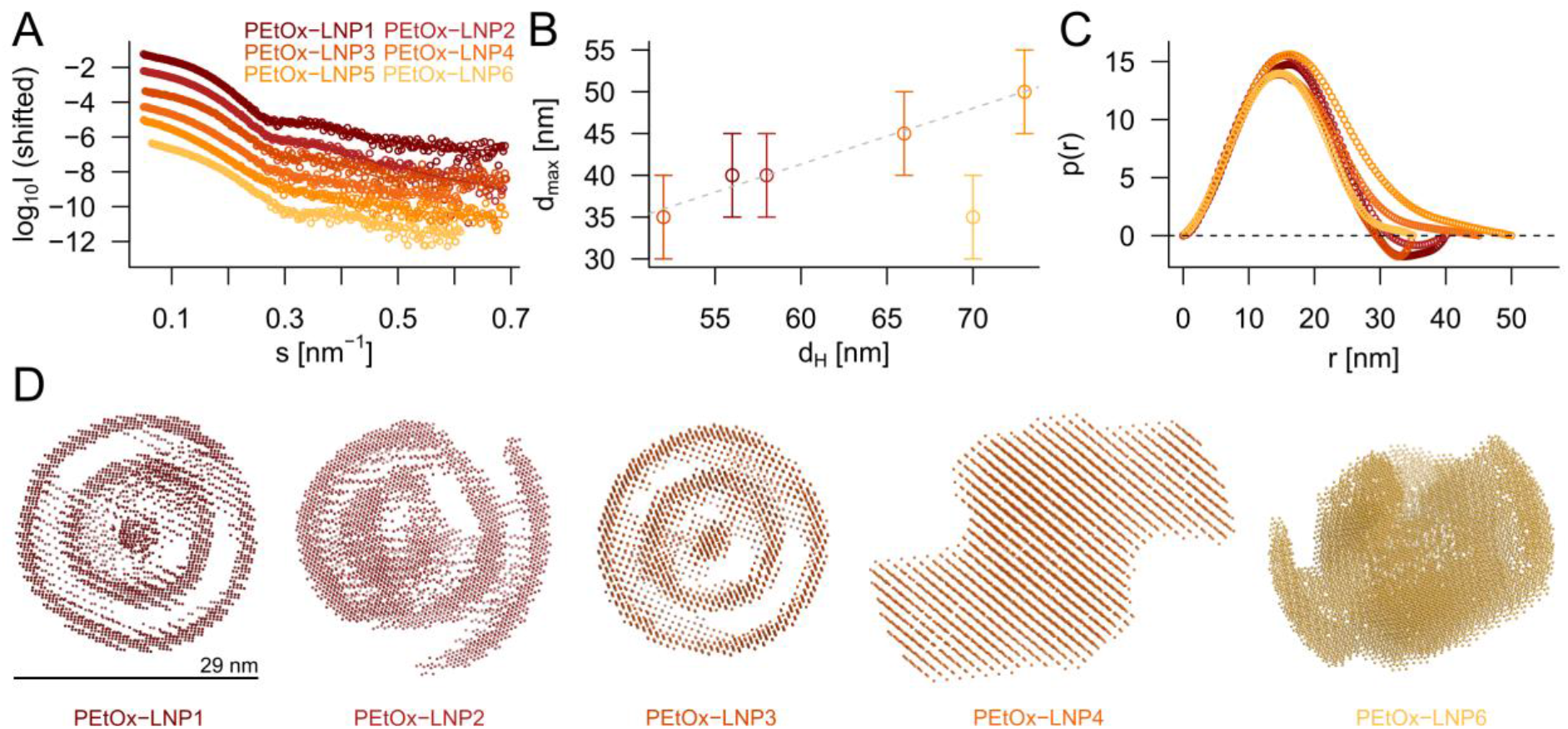
SAXS results for PEtOxylated LNPs. (A) Scattering curves of LNP samples, with fits shown as lines and intensities plotted as a function of momentum transfer. Log_10_I values for PEtOx-LNP2-6 were shifted by −1 to −5 for clarity. (B) Correlation between the maximum diameters determined by SAXS and d_H_ values as obtained from our microfluidic setup through DLS (colors according to (A), d_max_ shown as mean ± SD). (C) Distance distribution functions *p(r)* for all samples (colors according to (A)). (D) Orthoscopic view on *ab initio* bead models constructed by the program DAMMIN (rendered in PyMOL [63]). A scaling bar applying to all systems is provided below PEtOx-LNP1.

To obtain further structural information about the particles, the shape-determination program DAMMIN was employed [45]. This program uses simulated annealing to recover the overall low-resolution structure of a measured object from scattering data, represented as a bead model. Using this method, we were able to generate models for PEtOx-LNP1-4, as well as the unloaded variant PEtOx-LNP6 (**Figure 5D**). The models for the smaller particles, PEtOx-LNP1-3 (with higher N/P ratios than PEtOx-LNP4-5, **Table S4**), were similar to each other and provided a good fit to the low-angle portion of the data, revealing spherical particle shapes with concentric bead layers. Overall, across samples PEtOx-LNP1-3, we determined an average periodicity of about 6.4 nm between these layers, possibly corresponding to the distance between the most RNA-dense regions in the particle interior or individual hexagonal phases [53]. Interestingly, these bead models showed no occupancy above 30 nm, even though we defined a DAMMIN search volume with a diameter of 40 nm. This coincides with the negative *p(r)* excursion > 30 nm in these particles, as discussed above. The shapes of PEtOx-LNP4 and unloaded PEtOx-LNP6 were distinctly different, showing somewhat anisometric shapes without a well-developed internal structure. As SAXS averages the contributions from all scattering volumes in the sample, such shapes could arise either from an increase in polydispersity (**Table S4**) or the presence of different morphologies (e.g., lamellar and hexagonal phases being present simultaneously [53]).

The obtained data did not allow for the construction of an analogous bead model from the low-angle portion of the PEtOx-LNP5 containing larger-sized particles. However, here, the scattering data displayed (weak but noticeable) features at middle angles, pointing to internal order. A broad maximum *s*_*max*_ was observed around a momentum transfer of 1.3 nm^-1^ (**Figure S5**). Previous LNP studies also reported a Bragg peak around this area, being associated with repeated stackings of mRNA and lipid layers in the form of inverted hexagonal phases [36, 64–66]. Using PEAK [41, 42], the region of the scattering data around this peak were fitted by a mixture of Gaussian and Lorentzian functions. From this fit, we determined a *d*-spacing of the regular packing of about 5 nm, similar to previous measurements of PEGylated LNPs [36]. The distance between these regular structures was *L* = 13.5 nm. Therefore, each PEtOx-LNP5 particle contained about 2 to 3 correlated layers (**Table S7**).

Because the PEtOx chains are located on the outer surface of the LNPs, we further analyzed our samples for surface fractality using the Porod exponent. In this approach, the intensity at higher angles was represented by a power law *I*(*s*) ~ *s*^−χ^ + *const*., where the constant term *const*. was used to account for the scattering contribution from structural fluctuations. Porod exponents of χ = 3.8 to 4 (**Table S7**) indicated defined particles with sharp boundaries and relatively smooth surfaces. Previously, lower χ numbers were observed for polysarcosine-based LNP formulations, which suggested an increase in surface roughness compared to PEGylated systems [10]. We could not observe similar behavior for the herein-produced PEtOxylated LNPs.

### 3.3 Effects of altered polymer coatings on LNP biodistribution

To further link changes of surface characteristics within PEtOxylated LNPs to pharmacokinetic properties, we performed an *in vivo* barcoding study in mice. Here, the biodistribution of LNPs coated with PEtOx-lipids was compared to well-established PEGylated benchmarks, represented by the PEG-lipid (referred to as Comirnaty in the following section) and Spikevax formulations (the latter containing DMG-PEG2000 as PEGylated and SM-102 as ionizable lipid). We obtained well-defined particles with small sizes, low PDI, high EE, and near-zero ζ-potentials (**Table S9**). Additionally, prior to animal testing, we assessed the toxicity of the PEtOxylated LNPs and observed high cell viability at 200 ng/well, comparable to that of PEGylated variants (**Figure S6**). Each LNP was loaded with a FLuc mRNA variant and a single-stranded, uniquely identifiable DNA barcode. After intravenous injection, DNA barcode sequences associated with LNPs were isolated from tissues and quantified in a tissue-dependent manner using next-generation sequencing at 6 and 24 h (**Table S10-11**). For both time points, the highest abundance of absolute DNA barcodes was recovered for the Spikevax-based LNPs (**Figure S7**). Furthermore, in accordance with a deviation in the base lipid composition (50% SM-102 as ionizable lipid, instead of 46.3% ALC-0315), the Spikevax-based formulation exhibited a distinctly different biodistribution pattern, when compared to both PEGylated and PEtOxylated variants of the Comirnaty benchmark formulation.

To enable a detailed comparison of organ-specific LNP accumulation, we calculated the fraction of barcodes found within each organ based on the total amount of barcodes detected within the respective animal (**Figure 6A**). Relatively large fractions of LNPs associated with both the bone marrow and the whole blood at 6 h could correspond to systemically circulating particles prior to cellular uptake and endosomal/lysosomal processing, as well as an early distribution to highly perfused tissues. High amounts were also observed at this time point for the liver and the spleen, in accordance with the often-reported tropism of LNPs to these organs [67], as well as for the pancreas. At 24 h, the total amount of detectable barcodes decreased to 20 to 30% of the amounts recorded at 6 hours (**Table S11**), suggesting clearance of the injected nanoparticles within the investigated time frame. Compared to the 6-h-results, fractions within the bone marrow and the blood decreased, while the number of barcodes associated with the spleen increased, as seen most strikingly for the Spikevax formulation. Distribution to the brain, heart, or kidney was not observed for any formulation. Interestingly, at 6 h post-injection, PEtOxylated LNPs showed the highest fraction of barcodes across all LNP variants within the bone marrow (**Figure 6B**). No comparable significant differences between formulation types regarding any other organ, including between the two PEGylated benchmarks, were detected (**Figure S8A**). At 24 h, relative distributions across organs varied drastically among individual animals (**Figure 6A**, 24-h plot), resulting in no significant differences detectable between any formulation-organ combination (**Figure S8B**).

**Figure 6:**
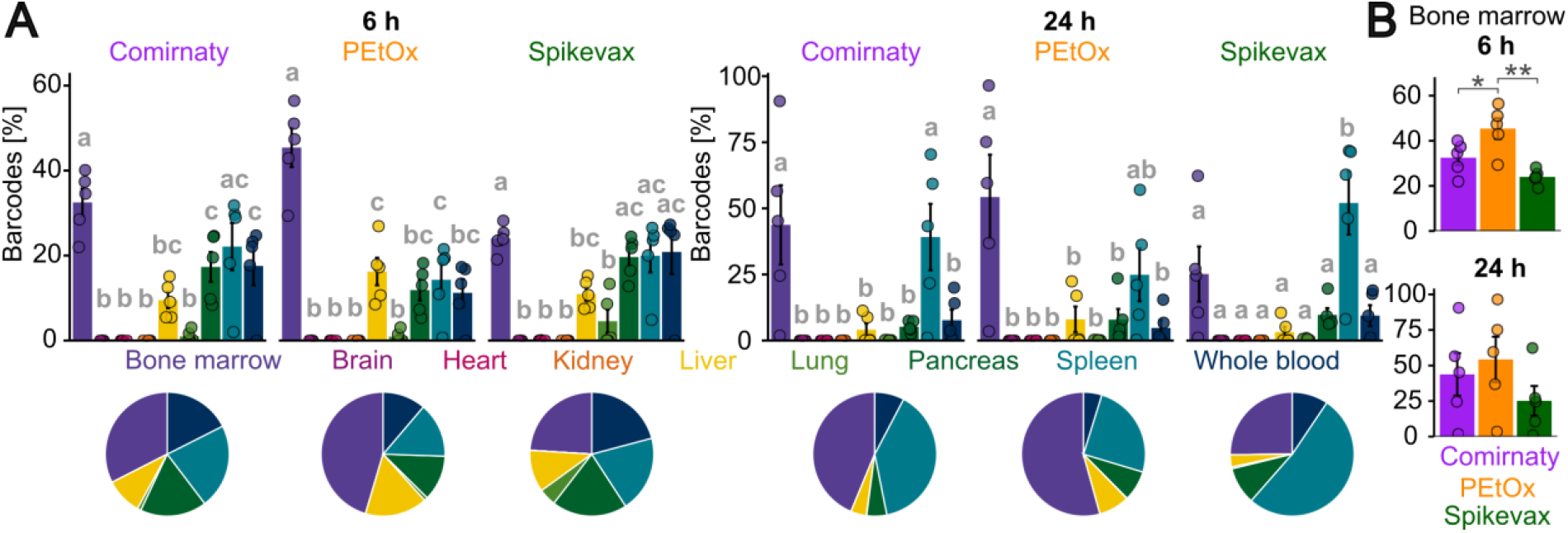
LNP biodistribution results. (A) Barplots and pie charts for the distribution of DNA barcodes 6 and 24 h post injection across all organs (mean ± SEM, n=5; significant differences (p < 0.05) indicated with a compact letter display (small letters a, b, and c), based on one-way ANOVA with post-hoc Tukey test, performed separately for each formulation and time point). Here, the fractions of barcodes found within an organ in relation to all recovered barcodes were computed for each animal separately, with individual values illustrated as points. (B) Fraction of barcodes found within the bone marrow after 6 and 24 h, compared between the three different LNP variants (mean ± SEM, n=5; significant differences indicated by stars based on one-way ANOVA with post-hoc Tukey test: * p < 0.05, ** p < 0.01, see also **Figure S8**).

The observed early-phase biodistribution shift towards the highly vascularized bone marrow upon a PEG-to-PEtOx exchange raises a point for discussion: As we did not observe a similar increase in the relative abundance of barcodes in whole blood when comparing PEtOxylated to PEGylated variants, the accumulation seen with excipient exchange is likely not a consequence of differences in terminal blood half-live. Instead, surface structural differences may account for the tropism of the PEtOxylated LNPs toward the bone marrow region. Particle adhesion to the fenestrated bone marrow sinusoidal endothelium at low velocity may be enhanced, thereby potentially increasing interactions with a wide array of scavenger receptors mediating uptake, such as stabilin-2 [68, 69]. Alternatively, the herein-reported surface structural changes could also shift the protein corona profile of particles, thereby altering the organ tropism of LNPs [70, 71]. To test our interpretation considering substantial variation across individual animals, we reanalyzed the data after an additional filtering step (excluding values outside the interquartile ranges), yielding qualitatively similar conclusions and suggesting that our interpretation is not due to an overinterpretation of extreme values (**Figure S9-11**). Here, a reduced fraction of barcodes in the whole blood 6 h post injection of PEtOxylated LNPs was additionally classified as significant (**Figure S11A**). Of note, DNA barcoding approaches provide information on the LNP biodistribution only and do not allow conclusions regarding functional delivery (protein expression from the delivered mRNA). Future work could probe the therapeutic potential resulting from such altered biodistribution profiles.

## 4 Conclusions

The self-regulating microfluidic machine, initially trained with PEGylated lipids, independently expanded its formulation capacities to POxylated lipids. Irrespective of the type of POx-lipid, the machine met preset targets faster for larger LNPs. PMeOxylated particles in the machine’s first attempt typically yielded larger diameters, similar to PEGylated lipids, whereas PEtOxylated LNPs initially yielded smaller diameters. Despite this polymer-dependent initial over- or underestimation of the predicted size, the machine always met the target. Smaller PEtOxylated LNPs (< 60 nm) had negative excursions in pair distribution functions *p(r)* as measured by SAXS, possibly pointing to a mixing of the outer cholesterol-rich monolayer with PEtOx chains. Concentric shapes suggested periodic structures, presumably relating to the individual mRNA layers. The PEtOxylated LNPs revealed a similar surface roughness as their PEGylated counterparts, in contrast to other previously reported alternatives such as polysarcosine [10]. *In vivo*, an increased fraction of PEtOxylated LNPs reached the bone marrow early after injection, compared to PEGylated counterparts. These results, derived from pairing self-regulated and automated LNP production with high-throughput methods for assessing biodistribution, underscore the importance of carefully selecting excipients to control biodistribution, with potential implications for therapeutic efficacy and adverse events. Furthermore, they demonstrate the potential of poly(2-oxazoline)s as alternative stealth lipids, particularly for organ-specific applications.

## CRediT authorship contribution statement

**Josef Kehrein:** Writing – original draft, Writing – review & editing, Visualization, Methodology, Investigation, Formal analysis, Data curation, Conceptualization; **Elena Reus:** Writing – review & editing, Visualization, Software, Methodology, Investigation, Formal analysis, Data curation, Conceptualization; **Caroline T. Holick:** Writing – original draft, Writing – review & editing, Methodology, Investigation, Formal analysis; **Sebastian Mummel:** Writing – original draft, Methodology, Investigation, Formal analysis; **Ilayda Ates:** Writing – original draft, Writing – review & editing, Investigation, Formal analysis; **Markus Hafke:** Formal analysis; **Jonas Käsbach:** Investigation, Formal analysis; **Christine Weber:** Writing – review & editing, Formal analysis; Supervision; **Tessa Lühmann:** Supervision; **Stephanie Schubert:** Writing – review & editing, Supervision, Resources, Funding acquisition; **Jørgen Magnus:** Writing – review & editing, Supervision, Resources, Funding acquisition; **Florian A. Mann:** Writing – review & editing, Supervision, Resources, Funding acquisition; **Ulrich S. Schubert:** Writing – review & editing, Supervision, Resources, Funding acquisition; **Lorenz Meinel:** Writing – review & editing, Supervision, Resources, Funding acquisition, Conceptualization.

## Acknowledgements

This work was funded by the BMWE (Bundesministerium für Wirtschaft und Energie) within the BASE-Lipid consortium (grant numbers: 16LP401002, 16LP401003, 16LP401006, and 16LP401007). The MALDI-TOF mass spectrometer (rapifleX) was funded by the Thüringer Aufbaubank (TAB, grant number 2016IZN0009). The ToC graphic was created using Biorender.com. The authors gratefully acknowledge BIOSAXS for performing the SAXS measurements and analyses, and Evonik Operations GmbH for generously providing *N,N*-ditetradecylamine, ALC-0315, ALC-0159, and cholesterol used in this study. Furthermore, we thank Zuzanna Makowska and Filippos Klironomos from Nuvisan Innovation Campus Berlin for conducting the *in vivo* biodistribution studies and Vanessa Hamann (Bayer AG) for her contributions to the establishment of the barcoding methodology. We also thank David Sparfeld and Celina Nehring (Bayer AG) for performing the *in vitro* cell toxicity assays.

## Appendix A Supplementary data

Supplementary data to this article can be found online.

## References

[1] Y. Xu, C. Wang, F. Shen, Z. Dong, Y. Hao, Y. Chen, Z. Liu, L. Feng, Lipid-coated CaCO(3) nanoparticles as a versatile pH-responsive drug delivery platform to enable combined chemotherapy of breast cancer, ACS Appl. Bio Mater., 5 (2022) 1194–1201.

[2] T.T.H. Thi, E.J.A. Suys, J.S. Lee, D.H. Nguyen, K.D. Park, N.P. Truong, Lipid-based nanoparticles in the clinic and clinical trials: from cancer nanomedicine to COVID-19 vaccines, Vaccines (Basel), 9 (2021) 359.

[3] H. Zhang, S. Han, X. Zhang, J. Ma, L. Jin, Y. Wang, Y. Ma, T. Ma, F. Yu, G. Song, Novel lipid nanoparticle (LNP) delivery systems enabling the advancement of RNA therapeutics, Adv. Healthc. Mater., (2026) e05340.

[4] C. Hald Albertsen, J.A. Kulkarni, D. Witzigmann, M. Lind, K. Petersson, J.B. Simonsen, The role of lipid components in lipid nanoparticles for vaccines and gene therapy, Adv. Drug Del. Rev., 188 (2022) 114416.

[5] Y. Eygeris, M. Gupta, J. Kim, G. Sahay, Chemistry of lipid nanoparticles for RNA delivery, Acc. Chem. Res., 55 (2022) 2–12.

[6] Q. Cheng, T. Wei, L. Farbiak, L.T. Johnson, S.A. Dilliard, D.J. Siegwart, Selective organ targeting (SORT) nanoparticles for tissue-specific mRNA delivery and CRISPR-CAS gene editing, Nat. Nanotechnol., 15 (2020) 313–320.

[7] S. Patel, N. Ashwanikumar, E. Robinson, Y. Xia, C. Mihai, J.P. Griffith, 3rd, S. Hou, A.A. Esposito, T. Ketova, K. Welsher, J.L. Joyal, O. Almarsson, G. Sahay, Naturally-occurring cholesterol analogues in lipid nanoparticles induce polymorphic shape and enhance intracellular delivery of mRNA, Nat. Commun., 11 (2020) 983.

[8] Y. Xu, S. Ma, H. Cui, J. Chen, S. Xu, F. Gong, A. Golubovic, M. Zhou, K.C. Wang, A. Varley, R.X.Z. Lu, B. Wang, B. Li, AGILE platform: a deep learning powered approach to accelerate LNP development for mRNA delivery, Nat. Commun., 15 (2024) 6305.

[9] L. Zhang, B.Y.L. Seow, K.H. Bae, Y. Zhang, K.C. Liao, Y. Wan, Y.Y. Yang, Role of PEGylated lipid in lipid nanoparticle formulation for in vitro and in vivo delivery of mRNA vaccines, J. Control. Release, 380 (2025) 108–124.

[10] S.S. Nogueira, A. Schlegel, K. Maxeiner, B. Weber, M. Barz, M.A. Schroer, C.E. Blanchet, D. Svergun, S. Ramishetti, D. Peer, P. Langguth, U. Sahin, H. Haas, Polysarcosine-functionalized lipid nanoparticles for therapeutic mRNA delivery, ACS Appl. Nano Mater., 3 (2020) 10634–10645.

[11] K. Su, L. Shi, T. Sheng, X. Yan, L. Lin, C. Meng, S. Wu, Y. Chen, Y. Zhang, C. Wang, Z. Wang, J. Qiu, J. Zhao, T. Xu, Y. Ping, Z. Gu, S. Liu, Reformulating lipid nanoparticles for organ-targeted mRNA accumulation and translation, Nat. Commun., 15 (2024) 5659.

[12] M. Mehta, T.A. Bui, X. Yang, Y. Aksoy, E.M. Goldys, W. Deng, Lipid-based nanoparticles for drug/gene delivery: an overview of the production techniques and difficulties encountered in their industrial development, ACS Mater. Au., 3 (2023) 600–619.

[13] L. Gurba-Bryskiewicz, W. Maruszak, D.A. Smuga, K. Dubiel, M. Wieczorek, Quality by design (QbD) and design of experiments (DOE) as a strategy for tuning lipid nanoparticle formulations for RNA delivery, Biomedicines, 11 (2023) 2752.

[14] C. Devos, A. Udepurkar, P. Sagmeister, A.S. Hodlewsky, J. Chen, A. Hatas, N. Ostrovsky, M. Al-Jazrawe, J.I. Ren, A.Y. Liu, R.D. Braatz, A.S. Myerson, Manufacturing mRNA-loaded lipid nanoparticles with precise size and morphology control, ACS Nano, 19 (2025) 33991–34002.

[15] S.J. Shepherd, D. Issadore, M.J. Mitchell, Microfluidic formulation of nanoparticles for biomedical applications, Biomaterials, 274 (2021) 120826.

[16] E. Reus, J. Savinsky, S. Wennemaring, J. Kasbach, F. Kerkhoffs, J. Kehrein, S.B. Rauer, T. Lühmann, A.C. Adams, M. Wessling, J. Magnus, L. Meinel, Self-regulating microfluidic system for lipid nanoparticle production, J. Control. Release, 388 (2025) 114370.

[17] R. Hoogenboom, The future of poly(2-oxazoline)s, Eur. Polym. J., 179 (2022) 111521.

[18] M. Glassner, M. Vergaelen, R. Hoogenboom, Poly(2-oxazoline)s: A comprehensive overview of polymer structures and their physical properties, Polym. Int., 67 (2018) 32–45.

[19] P. Hamm, M.D. Driessen, N. Hauptstein, J. Kehrein, R. Worschech, P. Pouyan, R. Haag, U.S. Schubert, T.D. Muller, L. Meinel, T. Lühmann, Deciphering polymer interactions in bioconjugates with different architectures by structural analysis via time-resolved limited proteolysis mass spectrometry, Angew. Chem. Int. Ed., 64 (2025) e202415354.

[20] N. Hauptstein, P. Pouyan, K. Wittwer, G. Cinar, O. Scherf-Clavel, M. Raschig, K. Licha, T. Lühmann, I. Nischang, U.S. Schubert, C.K. Pfaller, R. Haag, L. Meinel, Polymer selection impacts the pharmaceutical profile of site-specifically conjugated Interferon-alpha2a, J. Control. Release, 348 (2022) 881–892.

[21] N. Hauptstein, L. Meinel, T. Lühmann, Bioconjugation strategies and clinical implications of Interferon-bioconjugates, Eur. J. Pharm. Biopharm., 172 (2022) 157–167.

[22] N. Hauptstein, M. Dirauf, K. Wittwer, G. Cinar, O. Siering, M. Raschig, T. Lühmann, O. Scherf-Clavel, B. Sawatsky, I. Nischang, U.S. Schubert, C.K. Pfaller, L. Meinel, PEtOxylated interferon-alpha2a bioconjugates addressing H1N1 influenza A virus infection, Biomacromolecules, 23 (2022) 3593–3601.

[23] D. Haas, N. Hauptstein, M. Dirauf, M.D. Driessen, M. Ruopp, U.S. Schubert, T. Lühmann, L. Meinel, Chemo-enzymatic PEGylation/POxylation of murine interleukin-4, Bioconjug. Chem., 33 (2022) 97–104.

[24] N. Hauptstein, P. Pouyan, J. Kehrein, M. Dirauf, M.D. Driessen, M. Raschig, K. Licha, M. Gottschaldt, U.S. Schubert, R. Haag, L. Meinel, C. Sotriffer, T. Lühmann, Molecular insights into site-specific interferon-alpha2a bioconjugates originated from PEG, LPG, and PEtOx, Biomacromolecules, 22 (2021) 4521–4534.

[25] T. Lühmann, M. Schmidt, M.N. Leiske, V. Spieler, T.C. Majdanski, M. Grube, M. Hartlieb, I. Nischang, S. Schubert, U.S. Schubert, L. Meinel, Site-specific POxylation of interleukin-4, ACS Biomater. Sci. Eng., 3 (2017) 304–312.

[26] J.F.R. Van Guyse, S. Abbasi, K. Toh, Z. Nagorna, J. Li, A. Dirisala, S. Quader, S. Uchida, K. Kataoka, Facile generation of heterotelechelic poly(2-oxazoline)s towards accelerated exploration of poly(2-oxazoline)-based nanomedicine, Angew. Chem. Int. Ed., 63 (2024) e202404972.

[27] F.T. Kaps, A.-L. Ziegler, P. Fritsche, E. Takmakova, A. Kerr, S. Boye, A. Lederer, R. Luxenhofer, Electron-deficient alkyne lipids enable efficient synthesis of comparable polymer lipids via copper-free azide-alkyne cycloaddition, Angew. Chem. Int. Ed., 64 (2025) e202501262.

[28] X. He, T.J. Payne, A. Takanashi, Y. Fang, S.D. Kerai, J.P. Morrow, H. Al-Wassiti, C.W. Pouton, K. Kempe, Tailored monoacyl poly(2-oxazoline)- and poly(2-oxazine)-lipids as PEG-lipid alternatives for stabilization and delivery of mRNA-lipid nanoparticles, Biomacromolecules, 25 (2024) 4591–4603.

[29] A. Sanchez, D. Loughrey, E.S. Echeverri, S.G. Huayamares, A. Radmand, K. Paunovska, M. Hatit, K.E. Tiegreen, P.J. Santangelo, J.E. Dahlman, Substituting poly(ethylene glycol) lipids with poly(2-ethyl-2-oxazoline) lipids improves lipid nanoparticle repeat dosing, Adv. Healthc. Mater., 13 (2024) e2304033.

[30] B. Golba, Z. Zhong, M. Romio, R. Almey, D. Deforce, M. Dhaenens, N.N. Sanders, E.M. Benetti, B.G. De Geest, Cyclic poly(2-methyl-2-oxazoline)-lipid conjugates are good alternatives to poly(ethylene glycol)-lipids for lipid manoparticle mRNA formulation, Biomacromolecules, 26 (2025) 1816–1825.

[31] R.W. Moreadith, W. Kim, K. Smith, K. Yoon, R. Weimer, Z. Fang, Overcoming the PEG dilemma with poly(2-ethyl-2-oxazoline) lipids in lipid nanoparticle formulations, Eur. Polym. J., 241 (2025) 114392.

[32] M. Li, S. Ma, M. Shen, R. Wan, J.a. Long, Y. Jiang, H. Liang, Z. Guo, X. Ma, W. Song, Poly(2-oxazoline) decorated lipid nanoparticles for robust mRNA delivery in the presence of pre-existing anti-PEG antibodies, Sci. China Mater., (2026) 1–10.

[33] C.T. Holick, T. Klein, C. Mehnert, F. Adermann, I. Anufriev, M. Streiber, L. Harder, A. Traeger, S. Hoeppener, C. Franke, I. Nischang, S. Schubert, U.S. Schubert, Poly(2-ethyl-2-oxazoline) (POx) as poly(ethylene glycol) (PEG)-lipid substitute for lipid nanoparticle formulations, Small, 21 (2025) e2411354.

[34] J.T. Huckaby, T.M. Jacobs, Z. Li, R.J. Perna, A. Wang, N.I. Nicely, S.K. Lai, Structure of an anti-PEG antibody reveals an open ring that captures highly flexible PEG polymers, Commun. Chem., 3 (2020) 124.

[35] M.F.S. Deuker, V. Mailander, S. Morsbach, K. Landfester, Anti-PEG antibodies enriched in the protein corona of PEGylated nanocarriers impact the cell uptake, Nanoscale Horiz., 8 (2023) 1377–1385.

[36] B. Kolb, M. Graewert, R. Drexel, F. Meier, J. Raab, C. Wilhelmy, T. Nawroth, D. Soloviov, H. Haas, P. Langguth, Advanced quality and comparability assessment of mRNA-loaded lipid nanoparticles: absolute size distribution profiles and structure from AF4-coupled light and X-ray scattering measurements, Anal. Chem., 2026 (2026) 5271–5283.

[37] J. Schroeder, J. Ismail, C.T. Holick, J. Jungwirth, L. Klement, S. Hoeppener, C. Kosan, M. Schmidtke, B. Löffler, C. Weber, U.S. Schubert, C. Hoffmann, S. Schubert, C. Ehrhardt, Advancing influenza virus treatment: in vitro and ex vivo studies of PI3K inhibitor-loaded lipid nanoparticles, Mater. Today Bio, 35 (2025) 102587.

[38] F. Pedregosa, G. Varoquaux, A. Gramfort, V. Michel, B. Thirion, O. Grisel, M. Blondel, P. Prettenhofer, R. Weiss, V. Dubourg, J. Vanderplas, A. Passos, D. Cournapeau, M. Brucher, M. Perrot, E. Duchesnay, Scikit-learn: machine Learning in python, J. Mach. Learn. Res., 12 (2011) 2825–2830.

[39] N.R. Hajizadeh, D. Franke, D.I. Svergun, Integrated beamline control and data acquisition for small-angle X-ray scattering at the P12 BioSAXS beamline at PETRAIII storage ring DESY, J. Synchrotron Rad., 25 (2018) 906–914.

[40] C.E. Blanchet, A. Spilotros, F. Schwemmer, M.A. Graewert, A. Kikhney, C.M. Jeffries, D. Franke, D. Mark, R. Zengerle, F. Cipriani, S. Fiedler, M. Roessle, D.I. Svergun, Versatile sample environments and automation for biological solution X-ray scattering experiments at the P12 beamline (PETRA III, DESY), J. Appl. Crystallogr., 48 (2015) 431–443.

[41] K. Manalastas-Cantos, P.V. Konarev, N.R. Hajizadeh, A.G. Kikhney, M.V. Petoukhov, D.S. Molodenskiy, A. Panjkovich, H.D.T. Mertens, A. Gruzinov, C. Borges, C.M. Jeffries, D.I. Svergun, D. Franke, ATSAS 3.0: expanded functionality and new tools for small-angle scattering data analysis, J. Appl. Crystallogr., 54 (2021) 343–355.

[42] P.V. Konarev, V.V. Volkov, A.V. Sokolova, M.H.J. Koch, D.I. Svergun, PRIMUS: a Windows PC-based system for small-angle scattering data analysis, J. Appl. Crystallogr., 36 (2003) 1277–1282.

[43] A. Guinier, La diffraction des rayons X aux très petits angles : application à l’étude de phénomènes ultramicroscopiques, Ann. Phys. (N.Y.), 11 (1939) 161–237.

[44] D.I. Svergun, Determination of the regularization parameter in indirect-transform methods using perceptual criteria, J. Appl. Crystallogr., 25 (1992) 495–503.

[45] D.I. Svergun, Restoring low resolution structure of biological macromolecules from solution scattering using simulated annealing, Biophys. J., 76 (1999) 2879–2886.

[46] J.E. Dahlman, K.J. Kauffman, Y. Xing, T.E. Shaw, F.F. Mir, C.C. Dlott, R. Langer, D.G. Anderson, E.T. Wang, Barcoded nanoparticles for high throughput in vivo discovery of targeted therapeutics, Proc. Natl. Acad. Sci. U. S. A., 114 (2017) 2060–2065.

[47] C.D. Sago, S. Kalathoor, J.P. Fitzgerald, G.N. Lando, N. Djeddar, A.V. Bryksin, J.E. Dahlman, Barcoding chemical modifications into nucleic acids improves drug stability in vivo, J. Mater. Chem. B, 6 (2018) 7197–7203.

[48] M.P. Lokugamage, C.D. Sago, J.E. Dahlman, Testing thousands of nanoparticles in vivo using DNA barcodes, Curr. Opin. Biomed. Eng., 7 (2018) 1–8.

[49] M. Grube, M.N. Leiske, U.S. Schubert, I. Nischang, POx as an alternative to PEG? A hydrodynamic and light scattering study, Macromolecules, 51 (2018) 1905–1916.

[50] S. Li, Y. Hu, A. Li, J. Lin, K. Hsieh, Z. Schneiderman, P. Zhang, Y. Zhu, C. Qiu, E. Kokkoli, T.H. Wang, H.Q. Mao, Payload distribution and capacity of mRNA lipid nanoparticles, Nat. Commun., 13 (2022) 5561.

[51] C.B. Roces, G. Lou, N. Jain, S. Abraham, A. Thomas, G.W. Halbert, Y. Perrie, Manufacturing considerations for the development of lipid nanoparticles using microfluidics, Pharmaceutics, 12 (2020) 1095.

[52] M. Danaei, M. Dehghankhold, S. Ataei, F. Hasanzadeh Davarani, R. Javanmard, A. Dokhani, S. Khorasani, M.R. Mozafari, Impact of particle size and polydispersity index on the clinical applications of lipidic nanocarrier systems, Pharmaceutics, 10 (2018) 57.

[53] H. Li, P. Song, Y. Li, S. Tu, M. Mehmood, L. Chen, N. Li, Q. Tian, Mesoscopic structure of lipid nanoparticles studied by small-angle X-ray scattering: a spherical core-triple shell model analysis, Membranes (Basel), 15 (2025) 153.

[54] C. Wilhelmy, L. Uebbing, B. Kolb, M.A. Graewert, T. Nawroth, H. Haas, P. Langguth, Direct structural investigation of pH responsiveness in mRNA lipid nanoparticles: refining paradigms, J. Control. Release, 384 (2025) 113848.

[55] L. Liu, J.H. Kim, Z. Li, M. Sun, T. Northen, J. Tang, E. McIntosh, S. Karve, F. DeRosa, PEGylated lipid screening, composition optimization, and structure-activity relationship determination for lipid nanoparticle-mediated mRNA delivery, Nanoscale, 17 (2025) 11329–11344.

[56] D. Geißler, C. Gollwitzer, A. Sikora, C. Minelli, M. Krumrey, U. Resch-Genger, Effect of fluorescent staining on size measurements of polymeric nanoparticles using DLS and SAXS, Anal. Methods, 7 (2015) 9785–9790.

[57] B.J. Ree, J. Lee, Y. Satoh, K. Kwon, T. Isono, T. Satoh, M. Ree, A comparative study of dynamic light and X-ray scatterings on micelles of topological polymer amphiphiles, Polymers, 10 (2018) 1347.

[58] D. Urimi, M. Hellsing, N. Mahmoudi, C. Söderberg, R. Widenbring, L. Gedda, K. Edwards, T. Loftsson, N. Schipper, Structural characterization study of a lipid nanocapsule formulation intended for drug delivery applications using small-angle scattering techniques, Mol. Pharm., 19 (2022) 1068–1077.

[59] M.F.W. Trollmann, R.A. Bockmann, mRNA lipid nanoparticle phase transition, Biophys. J., 121 (2022) 3927–3939.

[60] J. Kehrein, R. Luxenhofer, A. Bunker, Moving beyond PEG: poly(2-oxazolines) and poly(2-oxazines) alter the surface properties of lipid nanoparticles, ChemRxiv (2025).

[61] M. Hammel, Y. Fan, A. Sarode, A.E. Byrnes, N. Zang, P. Kou, K. Nagapudi, D. Leung, C.C. Hoogenraad, T. Chen, C.W. Yen, G.L. Hura, Correlating the structure and gene silencing activity of oligonucleotide-loaded lipid nanoparticles using small-angle X-ray scattering, ACS Nano, 17 (2023) 11454–11465.

[62] F. Sebastiani, M. Yanez Arteta, M. Lerche, L. Porcar, C. Lang, R.A. Bragg, C.S. Elmore, V.R. Krishnamurthy, R.A. Russell, T. Darwish, H. Pichler, S. Waldie, M. Moulin, M. Haertlein, V.T. Forsyth, L. Lindfors, M. Cardenas, Apolipoprotein E binding drives structural and compositional rearrangement of mRNA-containing lipid nanoparticles, ACS Nano, 15 (2021) 6709–6722.

[63] Schrödinger LLC, The PyMOL molecular graphics system, version 3.1.0, 2015.

[64] L. Liu, J.-H. Kim, Z. Li, M. Sun, T. Northen, J. Tang, E. Mcintosh, S. Karve, F. DeRosa, PEGylated lipid screening, composition optimization, and structure–activity relationship determination for lipid nanoparticle-mediated mRNA delivery, Nanoscale, 17 (2025) 11329–11344.

[65] H. Yu, B.P. Dyett, C.J. Drummond, J. Zhai, Ionizable lipid nanoparticles for mRNA delivery: internal self-assembled inverse mesophase structure and endosomal escape, Acc. Chem. Res., 58 (2025) 3210–3222.

[66] C.R. Thorn, J.C. Hickey, Y.-C. Chi, M.M. Wang, K. Ng Huang, S.-J. Hong, Q. Zou, H.-M. Nguyen, R.H. Pak, P. Lim Soo, Alternative structural lipids impact mRNA-lipid nanoparticle morphology, stability, and activity, Mol. Pharm., 23 (2026) 209–223.

[67] M. Hosseini-Kharat, K.E. Bremmell, C.A. Prestidge, Why do lipid nanoparticles target the liver? Understanding of biodistribution and liver-specific tropism, Mol. Ther. Methods Clin. Dev., 33 (2025) 101436.

[68] P.H. Weigel, Discovery of the liver hyaluronan receptor for endocytosis (HARE) and its progressive emergence as the multi-ligand scavenger receptor stabilin-2, Biomolecules, 9 (2019) 454.

[69] H. Qian, S. Johansson, P. McCourt, B. Smedsrød, M. Ekblom, S. Johansson, Stabilins are expressed in bone marrow sinusoidal endothelial cells and mediate scavenging and cell adhesive functions, Biochem. Biophys. Res. Commun., 390 (2009) 883–886.

[70] M. Bros, L. Nuhn, J. Simon, L. Moll, V. Mailander, K. Landfester, S. Grabbe, The protein corona as a confounding variable of nanoparticle-mediated targeted vaccine delivery, Front. Immunol., 9 (2018) 1760.

[71] P.C. Ke, S. Lin, W.J. Parak, T.P. Davis, F. Caruso, A decade of the protein corona, ACS Nano, 11 (2017) 11773–11776.

