## Supplementary Information for "Exploring lipid nanoparticle design spaces using self-regulating microfluidic machines and multiplexed *in vivo* biodistribution"

### 1 Polymer synthesis

#### 1.1 Materials

4-(Ditetradecylamino)-4-oxobutanoic acid (SucDTDA) was prepared according to a modified literature protocol [1]. 2-Ethyl-2-oxazoline (EtOx, ≥99%, Sigma-Aldrich) was pre-dried over barium oxide (BaO, 90%, Acros) and distilled under inert conditions. 2-Methyl-2-oxazoline (MeOx, ≥99%, Thermo Fisher Scientific) was pre-dried over calcium hydride ($CaH_2$, 93%, Thermo Fisher Scientific) and distilled under inert conditions. Methyl tosylate (MeTos, 97%, Sigma-Aldrich) was dried over calcium hydride ($CaH_2$, Sigma-Aldrich) and distilled under reduced pressure. Acetic acid (AcOH, ACS, Reag. Ph. Eur., VWR), triethylamine ($NEt_3$, Sigma-Aldrich), sodium azide ($NaN_3$, ≥ 99%, Carl Roth), 0.5 M sodium methoxide (NaOMe) in methanol (MeOH, ≥ 99%, Sigma-Aldrich), triphenylphosphine ($PPh_3$, ≥ 99.5%, Carl Roth), *N*-hydroxysuccinimide (NHS, Sigma-Aldrich), 1-ethyl-3-(3-dimethylaminopropyl)carbodiimide (EDC, ≥ 97%, Sigma-Aldrich), 4-(*N,N*-dimethylamino)pyridine (DMAP, 99%, abcr), succinic anhydride (Sigma-Aldrich), 1-hydroxybenzotriazole monohydrate (HOBt × $H_2O$; ≥ 97% dry basis, Sigma-Aldrich), ethanol (99.5%, anhydrous, Fisher Scientific), and chloroform ($CHCl_3$, anhydrous, > 99%, Sigma-Aldrich) were used without further purification. Acetonitrile ($CH_3CN$), methanol (MeOH) and dichloromethane ($CH_2Cl_2$) were dried in a solvent purification system (SPS 800, MBRAUN). Technical grade diethylether ($Et_2O$) was used without further purification. *N,N*-Ditetradecylamine was kindly gifted by Evonik. Dialysis was performed using a Spectra/Por Biotech Cellulose Ester (CE) dialysis membrane with a molecular weight cut-off (MWCO) of 100 to 500 Da or 1000 Da (Carl Roth).

#### 1.2 Instrumentation

##### *1.2.1 Nuclear magnetic resonance (NMR) spectroscopy*

Proton NMR ($^1H$ NMR) spectra were measured using a Bruker AC 300 MHz spectrometer and a Bruker Avance NEO 300 MHz spectrometer with a 5 mm PA BBO 300S1 BBF-H-D-05Z probe head and a SampleJet robot for automated high throughput sample processing. The measurements were conducted at room temperature using $CDCl_3$ or $CD_2Cl_2$ as a solvent. The residual non-deuterated solvent signal was used as a reference. The spectra were baseline-corrected using the software SpinWorks 4 or Top Spin.

##### *1.2.2 Matrix assisted laser desorption / ionization time-of-flight mass spectrometry (MALDI TOF MS)*

MALDI TOF MS was measured on a rapifleX MALDI-TOF/TOF instrument from Bruker Daltonics equipped with a smartbeam™ 3D laser (355 nm wavelength). The spectra were measured in the positive reflector mode. For measurement of PEtOx samples, *trans*-2-[3-(4-*tert*-butylphenyl)-2-methyl-2-propenylidene]malononitrile (DCTB) was used as matrix, and sodium trifluoroacetate (NaTFA) was added as a doping salt. For measurement of PMeOx samples, α-cyano-4-hydroxycinnamic acid (CHCA) was used without a doping salt. The recording was conducted using the manufacturer's software flexControl 4.0. Evaluation and processing of the recorded spectra was done using the manufacturer's software flexAnalysis 4.0, including baseline subtraction and external calibration using a 5000 g $mol^{-1}$ poly(methyl methacrylate) (PMMA) standard from Polymer Standard Services (PSS).

##### *1.2.3 Size exclusion chromatography (SEC)*

SEC for PEtOx samples was measured on a Shimadzu system equipped with a CBM-20A system controller, a LC-10AD VP pump, a RID-10A refractive index detector and a SDV linear S column from PSS at 40 °C using $CHCl_3$:$NEt_3$:*iso*Propanol (94:4:2) as eluent at a flow rate of 1 mL/min. The calibration was made of polystyrene standards of narrow molar mass distribution (supplier PSS, $M_p$ = 370 to 128,000 g/mol). SEC for PMeOx samples was measured on an Agilent 1260 Infinity II system equipped with a G7110B pump, a G7162A refractive index detector and a guard GRAM 10 µm column, two GRAM 1000A 10 µm columns and a GRAM 30A 10 µm columns from Agilent at 50 °C using DMAc with 0.21 wt.% LiCl as eluent at a flow rate of 1 mL/min. The calibration was made of polystyrene standards of narrow molar mass distribution (supplier PSS, $M_p$ = 474 to 2,520,000 g/mol).

#### 1.3 Synthesis procedure

##### *1.3.1 PEtOx-lipid*

PEtOx-lipid was synthesized following a previously reported four-step procedure (**Scheme S1**) [2]. In brief, 2-ethyl-2-oxazoline was polymerized *via* cationic ring-opening polymerization (CROP). To introduce an acetate end group (PEtOx-OAc), the CROP was terminated with AcOH and $NEt_3$. Through post-polymerization modifications, the end group was first converted to a hydroxyl functionality (PEtOx-OH) by NaOMe-catalyzed transesterification, then to a carboxylic acid group yielding PEtOx-COOH by reaction with succinic anhydride. Finally, amidation of the latter with *N,N*-ditetradecylamine yielded the desired PEtOx-lipid. All species were characterized by means of $^{1}$H NMR spectroscopy, SEC, and MALDI TOF MS (**Figure S1-2**, **Table S1**).

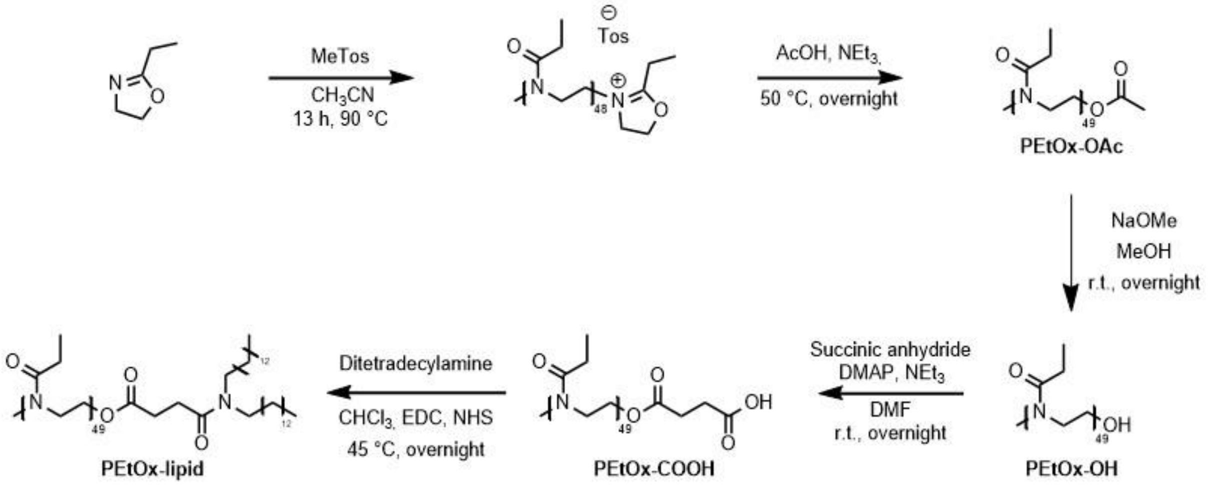


**Scheme S1:** Schematic representation of the PEtOx-lipid synthesis.

**PEtOx-OAc.** According to a [monomer] to [initiator] ratio of 50:1 and a monomer concentration of 4 mol/L, EtOx (20 g, 201.8 mmol, 50 eq.) and MeTos (751 mg, 4.04 mmol, 1 eq.) were dissolved in anhydrous $CH_3CN$ (30 mL). The reaction was stirred under reflux for 13 h and terminated by adding acetic acid (363.5 mg, 6.05 mmol, 1.5 eq.) and $NEt_3$ (816 mg, 8.07 mmol, 2 eq.). The reaction mixture was stirred at 50 °C overnight and was then diluted with $CHCl_3$, washed twice with aqueous $NaHCO_3$ solution and once with brine. The organic phase was dried over $Na_2SO_4$ and, after subsequent filtration, the solvent was removed under reduced pressure. The residue was dried at 40 °C *in vacuo*. The product was obtained as a white powder (16.4 g, 82%).

**PEtOx-OH.** PEtOx-OAc (16 g, 3.18 mmol, 1 eq.) was dissolved in anhydrous MeOH (107 mL), and NaOMe (0.5 M in MeOH, 640 µL, corresponding to 0.32 mmol (0.1 eq.) NaOMe) was added under vigorous stirring, which was continued at room temperature overnight. The solvent was removed under reduced pressure, and the residue was dissolved in chloroform. The mixture was washed twice with aqueous $NaHCO_3$

solution and brine. After drying the organic phase over $Na_2SO_4$ and filtration, the solvent was removed under reduced pressure. The residue was then dissolved in dichloromethane and precipitated in cold $Et_2O$ (−80 °C). The solid was dried *in vacuo*. The product was obtained as a white powder (14.7 g, 92%).

**PEtOx-COOH.** PEtOx-OH (14 g, 2.81 mmol, 1 eq.), succinic anhydride (425 mg, 4.25 mmol, 1.5 eq.) and DMAP (35 mg, 0.28 mmol, 0.1 eq.) were dissolved in anhydrous $CH_2Cl_2$ (35 mL). $NEt_3$ (426 mg, 4.21 mmol, 1.5 eq.) was added to the mixture, which was stirred overnight at room temperature. The solvent was removed under reduced pressure. The residue was dissolved in water and dialyzed against water (MWCO 0.1 to 0.5 kDa) for 3 days. The solution was lyophilized and the product was obtained as a white powder (8.17 g, 58%).

**PEtOx-lipid.** PEtOx-COOH (3.1 g, 0.60 mmol, 1 eq.), DMAP (8.4 mg, 0.06 mmol, 0.1 eq.), NHS (173 mg, 1.50 mmol, 2.5 eq.), and EDC (280 mg, 1.80 mmol, 3 eq.) were dissolved in anhydrous $CHCl_3$ (18 mL). The solution was stirred for 3 h at room temperature. *N,N*-Ditetradecylamine (493 mg, 1.20 mmol, 2 eq.) was added and the reaction mixture was stirred at 45 °C overnight. Afterwards, the mixture was diluted with $CHCl_3$, washed with aqueous $NaHCO_3$ solution and brine. After drying the organic phase over $Na_2SO_4$ and subsequent filtration, the solvent was removed under reduced pressure. The residue was dissolved in $CH_2Cl_2$ and precipitated in cold $Et_2O$ (−80 °C). The precipitate was dried *in vacuo*. The product was obtained as a white powder (1.63 g, 52%).

**Table S1:** Selected characterization data of PEtOx-lipid and its precursors.

| | $M_{n,\,theo.}$[a] [g mol$^{-1}$] | DP | DF[b] [%] | $M_{n,\,SEC}$[c] [g mol$^{-1}$] | $Đ_{SEC}$[c] | $M_{n,MALDI}$[d] [g mol$^{-1}$] | $Đ_{MALDI}$[d] |
|---|---|---|---|---|---|---|---|
| **PEtOx-OAc** | 4930 | 49 | 75 | 6070 | 1.07 | 4410 | 1.14 |
| **PEtOx-OH** | 4890 | 49 | 89 | 5690 | 1.07 | 4600 | 1.15 |
| **PEtOx-COOH** | 4990 | 49 | 65 | 5700 | 1.08 | 2110 | 1.66 |
| **PEtOx-lipid** | 5380 | 49 | 79 | 6850 | 1.05 | 4450 | 1.15 |

[a] Theoretical molar mass determined from $[M]_0/[I]_0$ and conversion. [b] Determined using $^1$H NMR spectroscopy. [c] Determined by SEC ($CHCl_3$, RI detection) using polystyrene calibration. [d] Determined by MALDI TOF MS (DCTB + NaTFA).

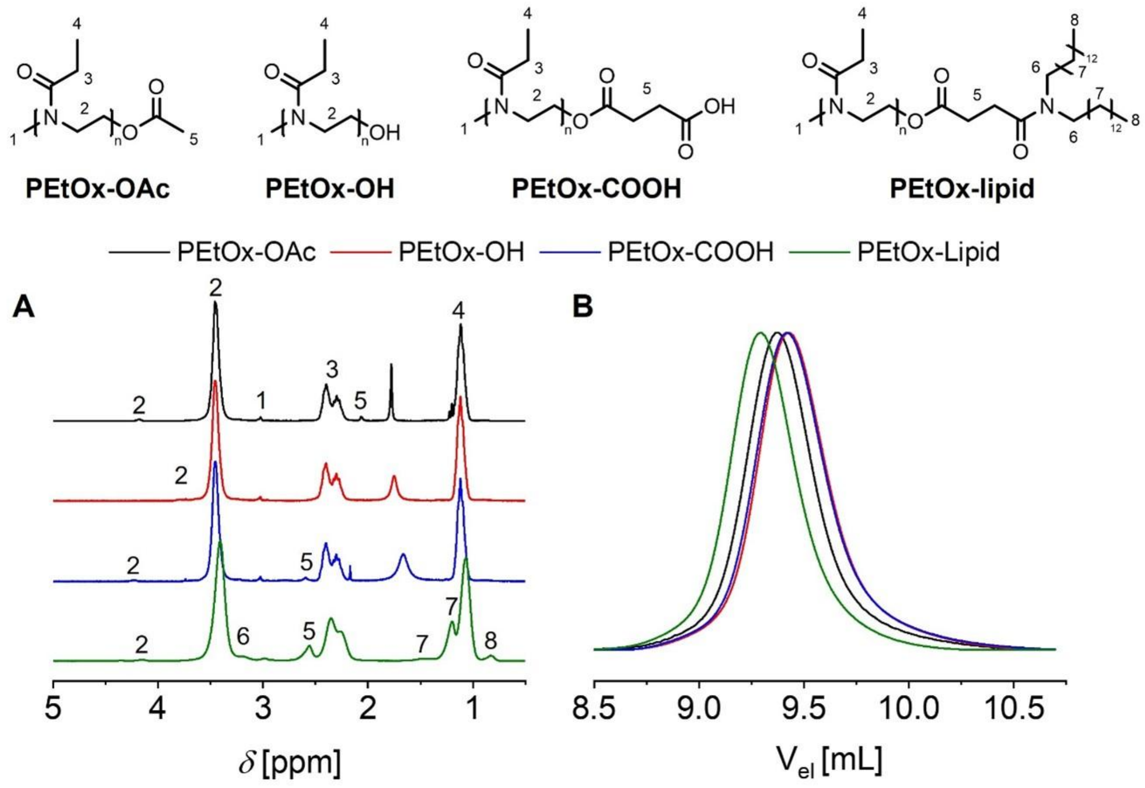


**Figure S1:** (A) $^{1}$H NMR spectra overlay of PEtOx-lipid and its precursors including peak assignment ($CDCl_3$, 300 MHz). (B) SEC overlay of PEtOx-lipid and its precursors (measured in $CHCl_3$, RI detection).

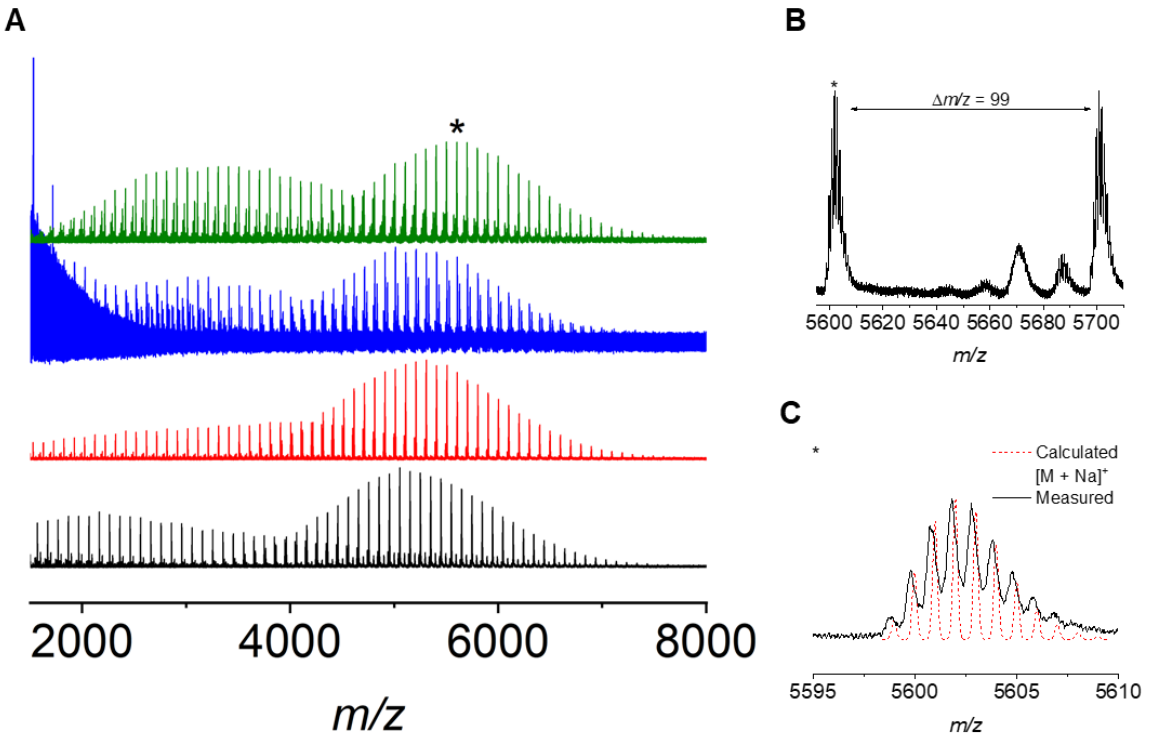


**Figure S2:** (A) Complete MALDI-TOF mass spectra of PEtOx-lipid and its precursors (DCTB, NaTFA); From bottom to top: PEtOx-OAc (black), PEtOx-OH (red), PEtOx-COOH (blue), PEtOx-lipid (green). (B) Display of the repeating unit of EtOx (Δ*m/z* = 99) in the spectrum of PEtOx-lipid. (C) Overlay of the measured and calculated isotopic patterns of the most abundant species (*) found for PEtOx-lipid.

##### *1.3.2 PMeOx-lipid*

PMeOx-lipid was synthesized in a 3-step procedure (**Scheme S2**). In brief, 2-methyl-2-oxazoline was polymerized *via* cationic ring-opening polymerization (CROP) and then terminated with sodium azide to introduce the azide end group (PMeOx-$N_3$). After reduction of the ω-end group yielding the primary amine (PMeOx-$NH_2$), coupling with 4-(ditetradecylamino)-4-oxobutanoic acid (SucDTDA) resulted in the desired PMeOx-lipid. All species were characterized by means of $^1$H NMR spectroscopy, SEC, and MALDI TOF MS (**Figure S3-4**, **Table S2**).

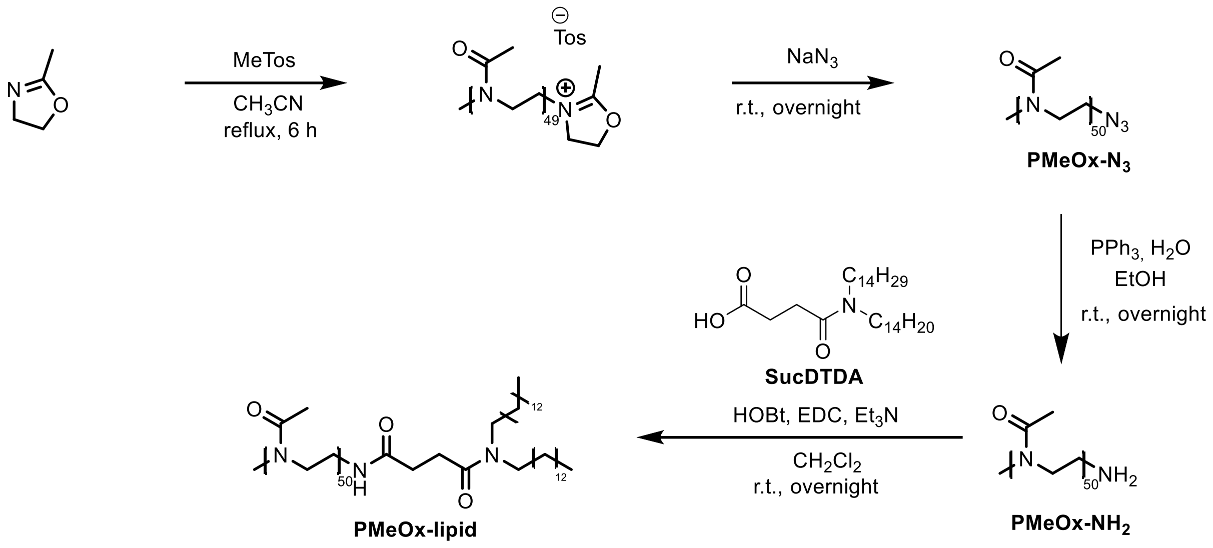


**Scheme S2:** Schematic representation of the PMeOx-lipid synthesis.

**PMeOx-$N_3$.** With a [monomer] to [initiator] ratio of 50:1, MeOx (300.0 g, 3.53 mol, 50 eq.) and MeTos (13.1 g, 0.07 mol, 1 eq.) were dissolved in anhydrous $CH_3CN$ (572 mL). The reaction mixture was stirred under reflux to achieve near quantitative monomer conversion, and subsequently terminated by adding sodium azide (9.2 g, 0.141 mol, 2 eq.). The reaction mixture was stirred at room temperature overnight. Subsequent to filtration, the solvent was removed under reduced pressure, and the residue was dried *in vacuo*. The product was obtained as a white powder (312 g, quantitative yield).

**PMeOx-$NH_2$.** Under an argon atmosphere, PMeOx-$N_3$ (130.0 g, 301 mmol, 1.00 eq.) and triphenylphosphine (262.3 g, 317 mmol, 1.05 eq.) were dissolved in anhydrous ethanol (1300 mL), and the solution was stirred at room temperature. After completion,

1 eq. water (540 µL, 301 mmol, 1.00 eq.) was added, and the mixture was stirred at room temperature until completion of the end group transformation. The resulting suspension was filtered, and the solvent was removed under reduced pressure. The residue was dried *in vacuo* and obtained as a white powder (138 g, quantitative yield).

**PMeOx-lipid.** PMeOx-$NH_2$ (8.0 g, 1.87 mmol, 1.0 eq.), SucDTDA (1.3 g, 1.87 mmol, 1.0 eq.), EDC (377 mg, 2.43 mmol, 1.30 eq.), HOBt × $H_2O$ (328 mg, 2.43 mmol, 1.30 eq.), and $NEt_3$ (0.37 ml, 2.43 mmol, 1.30 eq.) were dissolved in anhydrous $CH_2Cl_2$ (180 mL) and stirred at room temperature. After completion of the reaction, the product was precipitated from diethyl ether, dissolved in water, and dialyzed against water (MWCO 1.0 kDa). After lyophilization, a white powder was obtained (5.12 g, 57%).

**Table S2:** Selected characterization data of PMeOx-lipid and its precursors.

| | **$M_{n,\ theo.}$[a] [g mol$^{-1}$]** | **DP** | **DF[b] [%]** | **$M_{n,\ SEC}$[c] [g mol$^{-1}$]** | **$Đ_{SEC}$[c]** | **$M_{n,\ MALDI}$[d] [g mol$^{-1}$]** |
|---|---|---|---|---|---|---|
| **PMeOx-$N_3$** | 4310 | 50 | - | 7170 | 1.24 | 4900 |
| **PMeOx-$NH_2$** | 4280 | 50 | - | 6920 | 1.26 | 4570 |
| **PMeOx-lipid** | 4780 | 50 | 92 | 8720 | 1.18 | 4970 |

[a] Theoretical molar mass determined from $[M]_0/[I]_0$ and conversion. [b] Determined using $^1H$ NMR spectroscopy. [c] Determined by SEC using polystyrene calibration. [d] Determined by MALDI TOF MS (CHCA).

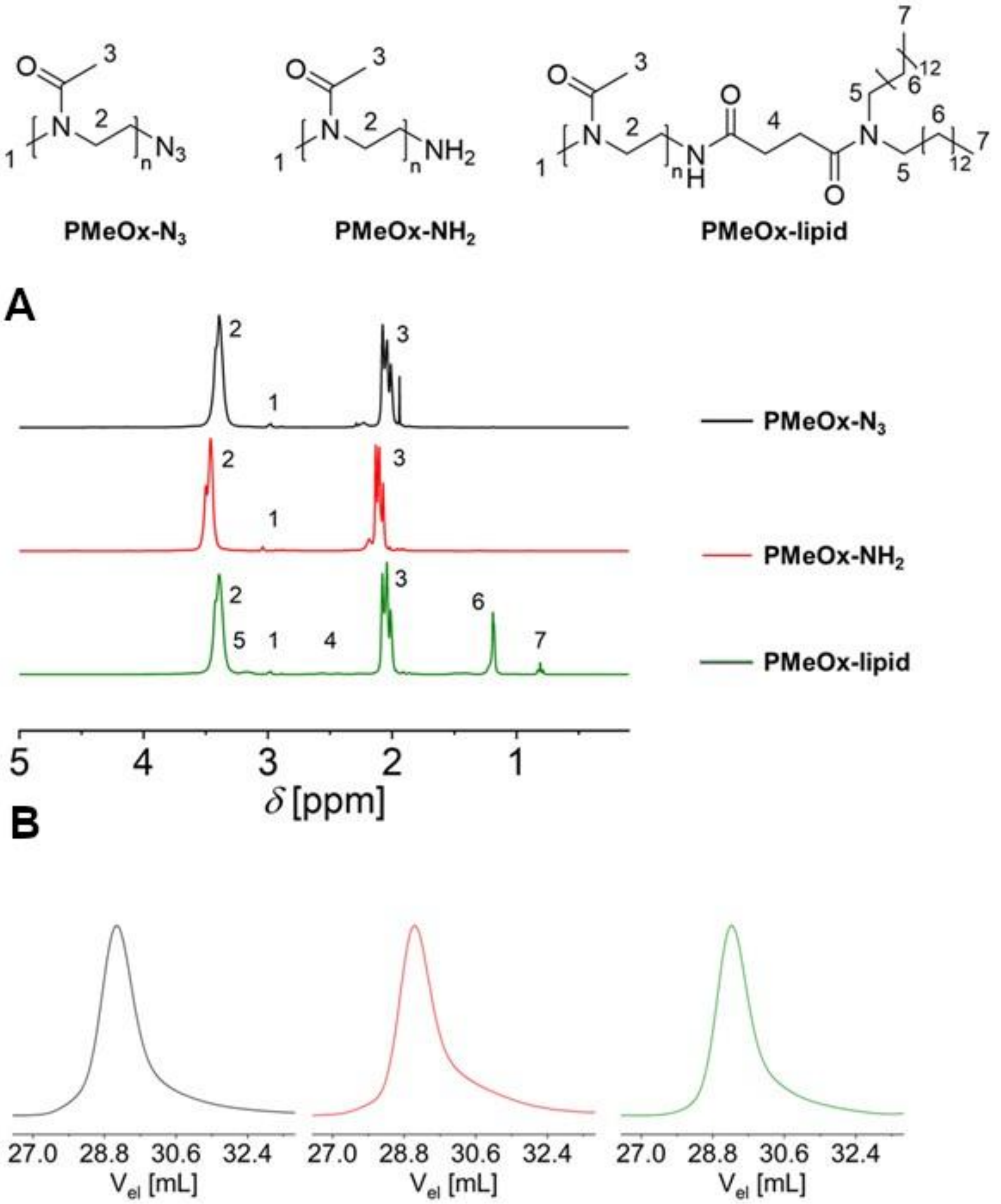


**Figure S3:** (A) $^1$H NMR spectra overlay of PMeOx-lipid and its precursors including peak assignment ($CDCl_3$ or $CD_2Cl_2$, 300 MHz). (B) SEC elugrams of PMeOx-lipid and its precursors (measured in DMAc, RI detection).

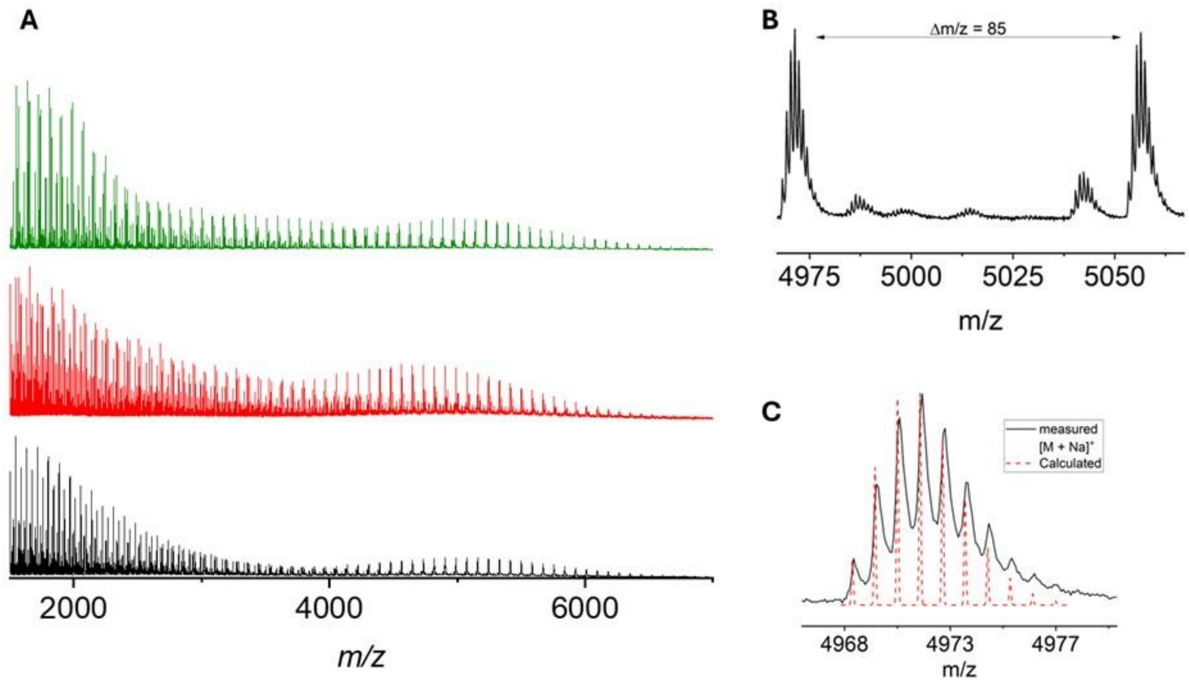


**Figure S4:** (A) Complete MALDI TOF mass spectra of PMeOx-lipid and its precursors (CHCA). From bottom to top: PMeOx-$N_3$ (black), PMeOx-$NH_2$ (red) and PMeOx-lipid (green). (B) Display of the repeating unit of MeOx ($\Delta m/z$ = 85) in the spectrum of PMeOx-lipid. (C) Overlay of the measured and calculated isotopic patterns of the most abundant species (*) found for PMeOx-lipid.

### 2 LNP formulation

**Table S3: Formulation parameters.** The table lists the allowed settings selectable within our microfluidic setup for the first run by our prediction model. Certain individual combinations were excluded due to pressure limits within our setup.

| Parameter | Allowed initial values |
|---|---|
| **RNA [mM]** | 0.1, 0.3, 0.5 |
| **Lipids [mM]** | 5, 12.5, 20 |
| **FRR** | 2, 3, 4, 5, 6 |
| **TFR [ml/min]** | 2, 2.25, 2.5, 2.75, 3, 3.25, 3.5, 3.75, 4, 4.25, 4.5, 4.75, 5, 5.25, 5.5, 5.75, 6 |

**Table S4: PEtOxylated LNPs formulated for SAXS measurements.** Particles are listed with their mean size (Z-average hydrodynamic diameter) and PDI determined in-line. PEtOx-LNP6 corresponds to empty LNPs used for comparison between loaded and unloaded particles.

| Name | mRNA [mM] | Lipids [mM] | TFR | FRR | N/P | Size [nm] | PDI |
|---|---|---|---|---|---|---|---|
| **PEtOx-LNP1** | 0.1 | 20 | 4.75 | 5 | 18.52 | 56 | 0.091 |
| **PEtOx-LNP2** | 0.3 | 20 | 4.75 | 4 | 7.72 | 58 | 0.063 |
| **PEtOx-LNP3** | 0.1 | 12.5 | 3.75 | 6 | 9.65 | 52 | 0.082 |
| **PEtOx-LNP4** | 0.3 | 12.5 | 3.75 | 4 | 4.82 | 66 | 0.115 |
| **PEtOx-LNP5** | 0.3 | 12.5 | 3.25 | 4 | 4.82 | 73 | 0.133 |
| **PEtOx-LNP6** | 0 | 12.5 | 4 | 4 | N/A | 70 | 0.153 |

### 3 SAXS measurements

**Table S5:** Technical specifications of the P12 beamline used for SAXS analysis.

| **Source** | **Petra III U29 undulator, P12 BioSAXS beamline** |
|---|---|
| **Detector** | Hybrid photon detector Pilatus 6M |
| **Energy range** | 10 keV |
| **Maximum flux at sample** | $1.00*10^{12}$ photons per second |
| **Beam size at sample** | 250 um x 100 um |
| **s-range** | 0.02 – 7.2 $nm^{-1}$ |
| **Wavelength** | 0.124 nm |
| **Sample-to-detector distance** | 3 m |
| **Axis calibration standard** | silver behenate |
| **Beamstop position** | off-set |

### 4 Self-regulating mechanism results

**Table S6: Self-regulating mechanism and LNP characterization results.** The table lists the initially selected target sizes along the settings suggested by the random forest prediction model. In the following columns, the experimentally determined sizes are listed and, if the size did not reach the desired 10% tolerance threshold, further adjustments of the TFR by the self-regulating mechanism are indicated with an arrow. Runs not reaching the tolerance interval are marked red, those that do are highlighted in green. Additionally, percentage values are given with the target value as reference. Furthermore, for each production run, PDIs along with EE values (mean values in %) are listed. Cells are marked pink for instances above the desired interval, and cyan for those below.

| Target value and initial settings | Size after 1st run and TFR adjustment for 2nd run | 2nd run and TFR adjustment for 3rd run | Size after 3rd run and TFR adjustment for 4th run | Size after 4th run |
|---|---|---|---|---|
| **PEGylated LNPs** | | | | |
| 61 nm<br>RNA: 0.1 mM<br>Lipids: 12.5 mM<br>TFR: 4.75<br>FRR: 6<br>N/P: 9.65 | 71 nm<br>115.74%<br><br>PDI: 0.214<br>EE: 84.4%<br>→ TFR: 5.25 | 61 nm<br>100.30%<br><br>PDI: 0.190<br>EE: 84.5% | - | - |
| 62 nm<br>RNA: 0.1 mM<br>Lipids: 12.5 mM<br>TFR: 4.75<br>FRR: 4<br>N/P: 14.47 | 75 nm<br>120.52%<br><br>PDI: 0.205<br>EE: 89.5%<br>→ TFR: 5.75 | 61 nm<br>99.13%<br><br>PDI: 0.180<br>EE: 94.1% | - | - |
| 63 nm<br>RNA: 0.1 mM<br>Lipids: 20 mM<br>TFR: 4.75<br>FRR: 6<br>N/P: 15.43 | 68 nm<br>108.15%<br><br>PDI: 0.234<br>EE: 90.11% | - | - | - |
| 69 nm<br>RNA: 0.3 mM<br>Lipids: 20 mM<br>TFR: 4.75<br>FRR: 5<br>N/P: 6.17 | 69 nm<br>99.32%<br><br>PDI: 0.198<br>EE: 88.3% | - | - | - |
| 70 nm<br>RNA: 0.3 mM<br>Lipids: 12.5 mM<br>TFR: 4.75<br>FRR: 6<br>N/P: 3.21 | 68 nm<br>97.45%<br><br>PDI: 0.166<br>EE: 88.2% | - | - | - |
| 75 nm<br>RNA: 0.3 mM<br>Lipids: 20 mM<br>TFR: 4.25<br>FRR: 4<br>N/P: 7.72 | 85 nm<br>113.35%<br><br>PDI: 0.202<br>EE: 88.2%<br>→ TFR: 4.75 | 80 nm<br>106.96%<br><br>PDI: 0.264<br>EE: 90.7% | - | - |
| 89 nm<br>RNA: 0.5 mM<br>Lipids: 12.5 mM<br>TFR: 3.75<br>FRR: 6<br>N/P: 1.93 | 102 nm<br>115.04%<br><br>PDI: 0.145<br>EE: 85.6%<br>→ TFR: 4.25 | 97 nm<br>108.45%<br><br>PDI: 0.198<br>EE: 75.5% | - | - |
| 90 nm<br>RNA: 0.3 mM<br>Lipids: 12.5 mM<br>TFR: 3.25<br>FRR: 3<br>N/P: 6.43 | 85 nm<br>94.71%<br><br>PDI: 0.189<br>EE: 94.5% | - | - | - |
| 97 nm<br>RNA: 0.5 mM<br>Lipids: 20 mM<br>TFR: 2<br>FRR: 4<br>N/P: 4.63 | 94 nm<br>97.33%<br><br>PDI: 0.196<br>EE: 80.9% | - | - | - |

| | | | | |
|---|---|---|---|---|
| 104 nm<br>RNA: 0.3 mM<br>Lipids: 12.5 mM<br>TFR: 2<br>FRR: 6<br>N/P: 3.22 | 104 nm<br>100.14%<br><br>PDI: 0.166<br>EE: 74.9% | - | - | - |
| **PMeOxylated LNPs** | | | | |
| 65 nm<br>RNA: 0.1 mM<br>Lipids: 20 mM<br>TFR: 4.75<br>FRR: 5<br>N/P: 18.52 | 68 nm<br>104.39%<br><br>PDI: 0.216<br>EE: 79.8% | - | - | - |
| 71 nm<br>RNA: 0.1 mM<br>Lipids: 12.5 mM<br>TFR: 4.25<br>FRR: 6<br>N/P: 9.65 | 84 nm<br>118.36%<br><br>PDI: 0.201<br>EE: 75.0%<br>→ TFR: 4.75 | 82 nm<br>115.81%<br><br>PDI: 0.198<br>EE: 75.0%<br>→ TFR: 5.25 | 68 nm<br>96.28%<br><br>PDI: 0.208<br>EE: 79.0% | - |
| 73 nm<br>RNA: 0.3 mM<br>Lipids: 20 mM<br>TFR: 4.25<br>FRR: 6<br>N/P: 5.14 | 91 nm<br>124.35%<br><br>PDI: 0.197<br>EE: 78.9%<br>→ TFR: 5.25 | 75 nm<br>102.66%<br><br>PDI: 0.203<br>EE: 77.4% | - | - |
| 78 nm<br>RNA: 0.3 mM<br>Lipids: 12.5 mM<br>TFR: 3.75<br>FRR: 4<br>N/P: 4.82 | 97 nm<br>124.66%<br><br>PDI: 0.159<br>EE: 77.4%<br>→ TFR: 4.75 | 77 nm<br>98.30%<br><br>PDI: 0.203<br>EE: 77.5% | - | - |
| 83 nm<br>RNA: 0.1 mM<br>Lipids: 5 mM<br>TFR: 4.25<br>FRR: 5<br>N/P: 4.63 | 73 nm<br>87.43%<br><br>PDI: 0.155<br>EE: 72.9%<br>→ TFR: 3.75 | 76 nm<br>91.84%<br><br>PDI: 0.160<br>EE: 74.6% | - | - |
| 87 nm<br>RNA: 0.5 mM<br>Lipids: 20 mM<br>TFR: 3.75<br>FRR: 6<br>N/P: 3.09 | 83 nm<br>94.84%<br><br>PDI: 0.187<br>EE: 75.5% | - | - | - |
| 91 nm<br>RNA: 0.3 mM<br>Lipids: 5 mM<br>TFR: 4.75<br>FRR: 4<br>N/P: 1.93 | 73 nm<br>80.12%<br><br>PDI: 0.108<br>EE: 76.2%<br>→ TFR: 4.25 | 82 nm<br>90.03%<br><br>PDI: 0.143<br>EE: 74.2% | - | - |
| 96 nm<br>RNA: 0.3 mM<br>Lipids: 12.5 mM<br>TFR: 2<br>FRR: 5<br>N/P: 3.86 | 93 nm<br>96.46%<br><br>PDI: 0.170<br>EE: 66.2% | - | - | - |
| 97 nm<br>RNA: 0.5 mM<br>Lipids: 20 mM<br>TFR: 2<br>FRR: 4<br>N/P: 4.63 | 100 nm<br>102.96%<br><br>PDI: 0.161<br>EE: 70.6% | - | - | - |
| 101 nm<br>RNA: 0.3 mM<br>Lipids: 5 mM<br>TFR: 4.25<br>FRR: 3<br>N/P: 2.57 | 96 nm<br>95.06%<br><br>PDI: 0.257<br>EE: 82.4% | - | - | - |
| 105 nm<br>RNA: 0.5 mM<br>Lipids: 12.5 mM<br>TFR: 2<br>FRR: 6<br>N/P: 1.93 | 113 nm<br>107.74%<br><br>PDI: 0.181<br>EE: 61.8% | - | - | - |

| | | | | |
|---|---|---|---|---|
| 107 nm<br>RNA: 0.5 mM<br>Lipids: 5 mM<br>TFR: 2.75<br>FRR: 2<br>N/P: 2.32 | 128 nm<br>119.27%<br><br>PDI: 0.125<br>EE: 82.3%<br>→ TFR: 3.25 | 120 nm<br>111.96%<br><br>PDI: 0.090<br>EE: 82.3%<br>→ TFR: 3.75 | 124 nm<br>116.10%<br><br>0.078<br>EE: 76.6%<br>→ STOPPED | - |
| **PEtOxylated LNPs** | | | | |
| 65 nm<br>RNA: 0.1 mM<br>Lipids: 20 mM<br>TFR: 5.25<br>FRR: 5<br>N/P: 18.52 | 58 nm<br>89.15%<br><br>PDI: 0.108<br>EE: 78.0%<br>→ TFR: 4.75 | 56 nm<br>86.63%<br><br>PDI: 0.122<br>EE: 79.0%<br>→ FRR: 4.25 | 62 nm<br>95.33%<br><br>PDI: 0.148<br>EE: 77.7% | - |
| 66 nm<br>RNA: 0.3 mM<br>Lipids: 20 mM<br>TFR: 4.75<br>FRR: 4<br>N/P: 7.72 | 58 nm<br>87.87%<br><br>PDI: 0.064<br>EE: 75.8%<br>→ TFR: 4.25 | 61 nm<br>92.52%<br><br>PDI: 0.087<br>EE: 72.3% | - | - |
| 78 nm<br>RNA: 0.3 mM<br>Lipids: 12.5 mM<br>TFR: 3.75<br>FRR: 4<br>N/P: 4.82 | 66 nm<br>84.21%<br><br>PDI: 0.115<br>EE: 73.9%<br>→ TFR: 3.25 | 73 nm<br>93.69%<br><br>PDI: 0.133<br>EE: 71.2% | - | - |
| 83 nm<br>RNA: 0.1 mM<br>Lipids: 5 mM<br>TFR: 4.25<br>FRR: 5<br>N/P: 4.63 | 71 nm<br>86.37%<br><br>PDI: 0.255<br>EE: 67.0%<br>→ TFR: 3.75 | 72 nm<br>87.77%<br><br>PDI: 0.137<br>EE: 65.1%<br>→ TFR: 3.25 | 74 nm<br>89.41%<br><br>PDI: 0.204<br>EE: 55.9%<br>→ STOPPED | - |
| 91 nm<br>RNA: 0.1 mM<br>Lipids: 5 mM<br>TFR: 4.25<br>FRR: 5<br>N/P: 1.93 | 70 nm<br>76.69%<br><br>PDI: 0.125<br>EE: 64.3%<br>→ TFR: 3.75 | 72 nm<br>79.22%<br><br>PDI: 0.111<br>EE: 62.5%<br>→ TFR: 2.75 | 74 nm<br>80.83%<br><br>PDI: 0.119<br>EE: 57.6%<br>→ TFR: 2.25 | 90 nm<br>99.33%<br><br>PDI: 0.145<br>EE: 62.2% |
| 105 nm<br>RNA: 0.5 mM<br>Lipids: 12.5 mM<br>TFR: 2<br>FRR: 6<br>N/P: 1.93 | 95 nm<br>90.05%<br><br>PDI: 0.132<br>EE: 66.7% | - | - | - |
| 107 nm<br>RNA: 0.5 mM<br>Lipids: 5 mM<br>TFR: 2.75<br>FRR: 2<br>N/P: 2.32 | 110 nm<br>102.37%<br><br>PDI: 0.106<br>EE: 63.1% | - | - | - |
| 111 nm<br>RNA: 0.5 mM<br>Lipids: 5 mM<br>TFR: 5.75<br>FRR: 6<br>N/P: 0.77 | 116 nm<br>104.87%<br><br>PDI: 0.161<br>EE: 36.2% | - | - | - |
| 119 nm<br>RNA: 0.5 mM<br>Lipids: 5 mM<br>TFR: 3.75<br>FRR: 6<br>N/P: 0.77 | 155<br>130.26%<br><br>PDI: 0.191<br>EE: 40.2%<br>→ TFR: 4.75 | 131 nm<br>109.93%<br><br>PDI: 0.157<br>EE: 35.2% | - | - |

### 5 SAXS results

**Table S7: SAXS results.** Particles are listed according to names from **Table S4**.

| Name | $R_g$ [nm] | $d_{max}$ [nm] | $s_{max}$ [$nm^{-1}$] | Spacing [nm] | L [nm] | Porod exponent |
|---|---|---|---|---|---|---|
| **PEtOx-LNP1** | 10.5 ± 0.5 | 40 ± 5 | | | | 4 |
| **PEtOx-LNP2** | 11.5 ± 0.5 | 40 ± 5 | | | | 4 |
| **PEtOx-LNP3** | 10.9 ± 0.5 | 35 ± 5 | | | | 4 |
| **PEtOx-LNP4** | 13.0 ± 0.5 | 45 ± 5 | | | | 3.9 |
| **PEtOx-LNP5** | 14 ± 0.5 | 50 ± 5 | 1.27 | 5.0 | 13.5 | 3.8 |
| **PEtOx-LNP6** | 11.5 ± 0.5 | 35 ± 5 | | | | 4 |

**Table S8: Electron densities of individual lipid components and solvent.** Values were computed via CRYSOL [3], using PDB structures of each molecule.

| Compound | Electron density [e/Å³] |
|---|---|
| **Water** | 0.334 |
| **ALC-0159 (PEG-lipid)** | 0.367 |
| **PEtOx-lipid** | 0.377 |
| **ALC-0315** | 0.324 |
| **DSPC** | 0.348 |
| **Cholesterol** | 0.312 |

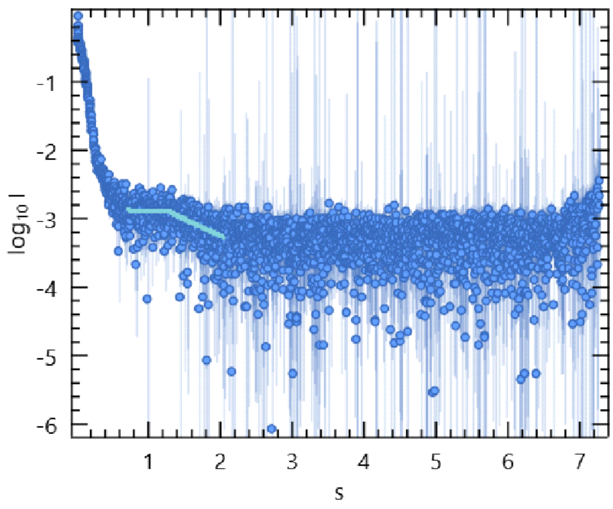


**Figure S5:** Scattering intensity of PEtOx-LNP5 illustrated with the fit in the region of the peak that was used for determining order parameters through PEAK.

### 6 Barcoding study

**Table S9: Characterization of LNPs.** Size, PDI, EE, and ζ values (mean values, n=3) for LNP batches that were produced by the NanoAssemblr™ system and used in our *in vivo* barcoding study.

| **Name** | **LNP polymer [mol%]** | **Ionizable lipid [mol%]** | **Phospholipid [mol%]** | **Cargo** | **Barcode** | **Size [nm]** | **PDI** | **EE [%]** | **ζ [mV]** |
|---|---|---|---|---|---|---|---|---|---|
| **Comirnaty** | ALC-0159 [1.6] | ALC-0315 [46.3] | DSPC [9.4] | mRNA-Fluc:ssDNA-bc_17 | CTCCTTCG | 62 | 0.04 | 91 | 0 |
| **PEtOx** | PEtOx-lipid [1.6] | ALC-0315 [46.3] | DSPC [9.4] | mRNA-Fluc:ssDNA-bc_05 | AGGCGCTA | 56 | 0.05 | 73 | -1.5 |
| **Spikevax** | DMG-PEG2000 [1.5] | SM102 [50] | DSPC [10] | mRNA-Fluc:ssDNA-bc_18 | ACGCTAGC | 86 | 0.04 | 95 | 0.2 |

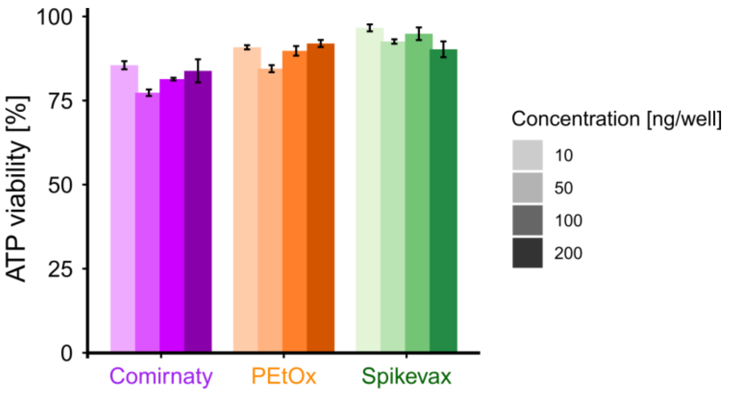


**Figure S6: ATP viability assay.** Results are shown per concentration and LNP type, normalized to negative (100%) and dead (0%) controls (mean ± SD, n=3).

**Table S10: Barcoding results.** Raw DNA barcode counts per measurement (color scaling applied column-wise from blue, low amounts, to red, high amounts). Animals were labelled with numbers 5 – 14, with IDs 5 – 9 investigated after 6 h, and IDs 10 – 14 after 24 h.

| Animal ID | Tissue | Time point | Comirnaty | PEtOx | Spikevax |
|---|---|---|---|---|---|
| Triplicate measurements of input barcodes to be injected (~ 80,000) | | | 83185 | 80388 | 78774 |
| | | | 82750 | 80070 | 78436 |
| | | | 88332 | 85856 | 84138 |
| 5 | Bone marrow | 6h | 29191 | 19138 | 73864 |
| 6 | Bone marrow | 6h | 25105 | 15015 | 75175 |
| 7 | Bone marrow | 6h | 32237 | 20600 | 82698 |
| 8 | Bone marrow | 6h | 36406 | 28063 | 82088 |
| 9 | Bone marrow | 6h | 30225 | 20780 | 81265 |
| 10 | Bone marrow | 24h | 11296 | 8844 | 28708 |
| 11 | Bone marrow | 24h | 315 | 216 | 1094 |
| 12 | Bone marrow | 24h | 14749 | 11958 | 29434 |
| 13 | Bone marrow | 24h | 6393 | 5078 | 11659 |
| 14 | Bone marrow | 24h | 14670 | 10998 | 33991 |
| 5 | Brain | 6h | 0 | 0 | 2 |
| 6 | Brain | 6h | 0 | 0 | 0 |
| 7 | Brain | 6h | 0 | 0 | 0 |
| 8 | Brain | 6h | 0 | 0 | 0 |
| 9 | Brain | 6h | 1 | 0 | 53 |
| 10 | Brain | 24h | 0 | 0 | 0 |
| 11 | Brain | 24h | 0 | 0 | 0 |
| 12 | Brain | 24h | 0 | 0 | 0 |
| 13 | Brain | 24h | 0 | 0 | 0 |
| 14 | Brain | 24h | 0 | 0 | 0 |
| 5 | Heart | 6h | 9 | 3 | 272 |
| 6 | Heart | 6h | 0 | 0 | 0 |
| 7 | Heart | 6h | 0 | 0 | 1 |
| 8 | Heart | 6h | 0 | 0 | 4 |
| 9 | Heart | 6h | 0 | 0 | 0 |
| 10 | Heart | 24h | 0 | 3 | 20 |
| 11 | Heart | 24h | 0 | 0 | 1 |
| 12 | Heart | 24h | 0 | 0 | 17 |
| 13 | Heart | 24h | 0 | 0 | 0 |
| 14 | Heart | 24h | 0 | 0 | 1 |
| 5 | Kidney | 6h | 0 | 0 | 0 |
| 6 | Kidney | 6h | 0 | 0 | 17 |
| 7 | Kidney | 6h | 1 | 0 | 19 |
| 8 | Kidney | 6h | 8 | 0 | 91 |
| 9 | Kidney | 6h | 1 | 0 | 56 |
| 10 | Kidney | 24h | 0 | 0 | 3 |
| 11 | Kidney | 24h | 0 | 0 | 1 |
| 12 | Kidney | 24h | 0 | 0 | 0 |
| 13 | Kidney | 24h | 0 | 0 | 2 |
| 14 | Kidney | 24h | 2 | 0 | 10 |
| 5 | Liver | 6h | 20051 | 17544 | 58977 |
| 6 | Liver | 6h | 6593 | 5474 | 31771 |
| 7 | Liver | 6h | 6150 | 5111 | 27395 |
| 8 | Liver | 6h | 12192 | 9816 | 46856 |
| 9 | Liver | 6h | 4129 | 3138 | 21476 |
| 10 | Liver | 24h | 2192 | 2517 | 6021 |
| 11 | Liver | 24h | 28 | 29 | 119 |
| 12 | Liver | 24h | 30 | 33 | 138 |
| 13 | Liver | 24h | 2933 | 3086 | 11515 |
| 14 | Liver | 24h | 4 | 3 | 11 |
| 5 | Lung | 6h | 177 | 65 | 2718 |
| 6 | Lung | 6h | 0 | 0 | 0 |
| 7 | Lung | 6h | 7 | 2 | 290 |
| 8 | Lung | 6h | 1590 | 781 | 29647 |
| 9 | Lung | 6h | 2314 | 1137 | 38424 |
| 10 | Lung | 24h | 44 | 26 | 1063 |
| 11 | Lung | 24h | 16 | 5 | 162 |
| 12 | Lung | 24h | 57 | 41 | 728 |
| 13 | Lung | 24h | 4 | 1 | 29 |
| 14 | Lung | 24h | 11 | 14 | 324 |
| 5 | Pancreas | 6h | 11089 | 3461 | 63664 |
| 6 | Pancreas | 6h | 7077 | 2312 | 37872 |
| 7 | Pancreas | 6h | 27842 | 8754 | 80333 |

| | | | | | |
|---|---|---|---|---|---|
| 8 | Pancreas | 6h | 23776 | 8553 | 74856 |
| 9 | Pancreas | 6h | 14715 | 4780 | 70389 |
| 10 | Pancreas | 24h | 636 | 343 | 7577 |
| 11 | Pancreas | 24h | 1273 | 1451 | 9112 |
| 12 | Pancreas | 24h | 1918 | 1260 | 7515 |
| 13 | Pancreas | 24h | 1099 | 581 | 6912 |
| 14 | Pancreas | 24h | 737 | 182 | 10703 |
| 5 | Spleen | 6h | 39434 | 14020 | 91387 |
| 6 | Spleen | 6h | 20986 | 6144 | 73371 |
| 7 | Spleen | 6h | 20586 | 5245 | 70441 |
| 8 | Spleen | 6h | 1947 | 384 | 16714 |
| 9 | Spleen | 6h | 23962 | 6987 | 76428 |
| 10 | Spleen | 24h | 10808 | 3117 | 73108 |
| 11 | Spleen | 24h | 12019 | 3518 | 71141 |
| 12 | Spleen | 24h | 5658 | 1557 | 50177 |
| 13 | Spleen | 24h | 15446 | 4995 | 78412 |
| 14 | Spleen | 24h | 172 | 40 | 4409 |
| 5 | Whole blood | 6h | 32861 | 10943 | 96167 |
| 6 | Whole blood | 6h | 12857 | 2745 | 78273 |
| 7 | Whole blood | 6h | 26287 | 8300 | 90623 |
| 8 | Whole blood | 6h | 21500 | 7369 | 93842 |
| 9 | Whole blood | 6h | 1 | 0 | 39 |
| 10 | Whole blood | 24h | 0 | 0 | 9 |
| 11 | Whole blood | 24h | 3418 | 953 | 17493 |
| 12 | Whole blood | 24h | 3670 | 1069 | 20588 |
| 13 | Whole blood | 24h | 173 | 42 | 1266 |
| 14 | Whole blood | 24h | 602 | 164 | 5100 |

**Table S11: Barcoding results.** Raw barcode amounts summarized per animal ID, and overall, for each time point.

| Animal | Comirnaty | PEtOx | Spikevax |
|---|---|---|---|
| **Animal 5** | 132812 | 65174 | 387051 |
| **Animal 6** | 72618 | 31690 | 296479 |
| **Animal 7** | 113110 | 48012 | 351800 |
| **Animal 8** | 97419 | 54966 | 344098 |
| **Animal 9** | 75348 | 36822 | 288130 |
| **Mean** | **98261** | **47333** | **333512** |
| **SD** | **22793** | **12101** | **36726** |
| **Animal 10** | 24976 | 14850 | 116509 |
| **Animal 11** | 17069 | 6172 | 99123 |
| **Animal 12** | 26082 | 15918 | 108597 |
| **Animal 13** | 26048 | 13783 | 109795 |
| **Animal 14** | 16198 | 11401 | 54549 |
| **Mean** | **22075** | **12425** | **97715** |
| **SD** | **4469** | **3466** | **22284** |
| **6h+24h sum** | **120336** | **59758** | **431226** |
| **mean(24h) / mean(6h) * 100%** | **22.47%** | **26.25%** | **29.30%** |

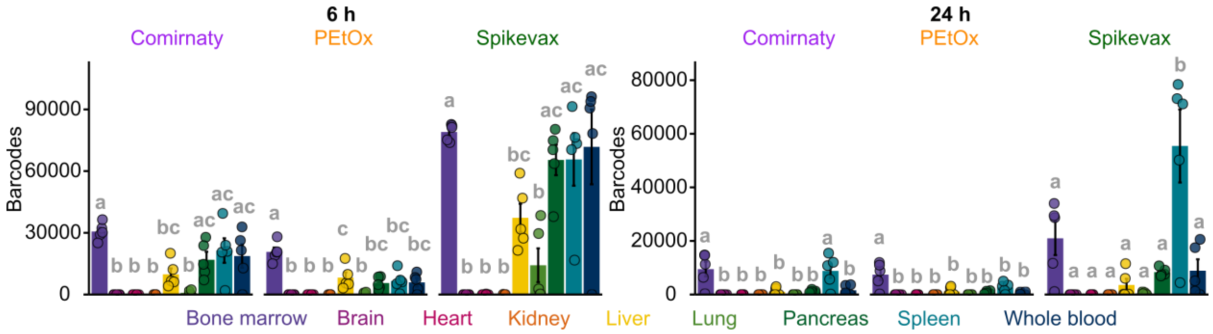


**Figure S7: Absolute DNA barcode amounts.** Results are shown per animal (points) for each organ separately (mean ± SEM, n=5; significant differences ($p < 0.05$) indicated with a compact letter display (small letters a, b, and c), based on one-way ANOVA with post-hoc Tukey test, performed separately for each formulation and time point).

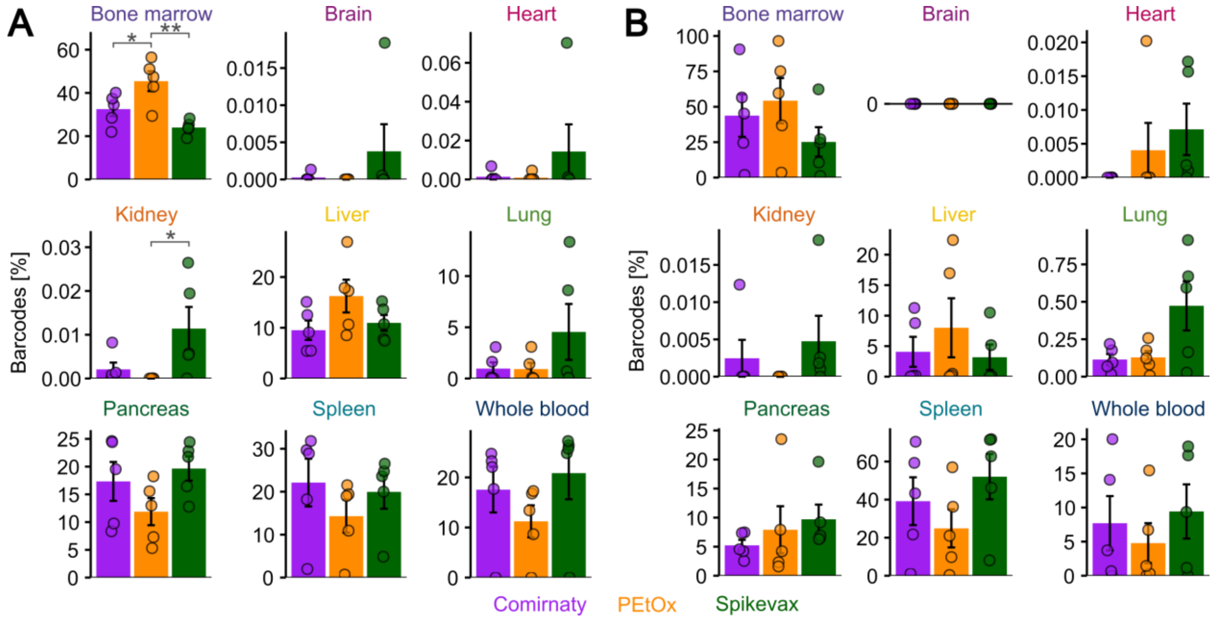


**Figure S8: Organ-dependent DNA barcoding results.** Fractions of barcodes found in specific organs are compared across formulations, with ALC-0159 results shown in purple, PEtOx results in orange, and Spikevax results in green (mean ± SEM, n=5, data according to **Figure 6A**, significant differences are indicated based on one-way ANOVA with post-hoc Tukey test: * $p < 0.05$, ** $p < 0.01$). (A) Results for 6 h. (B) Results for 24 h.

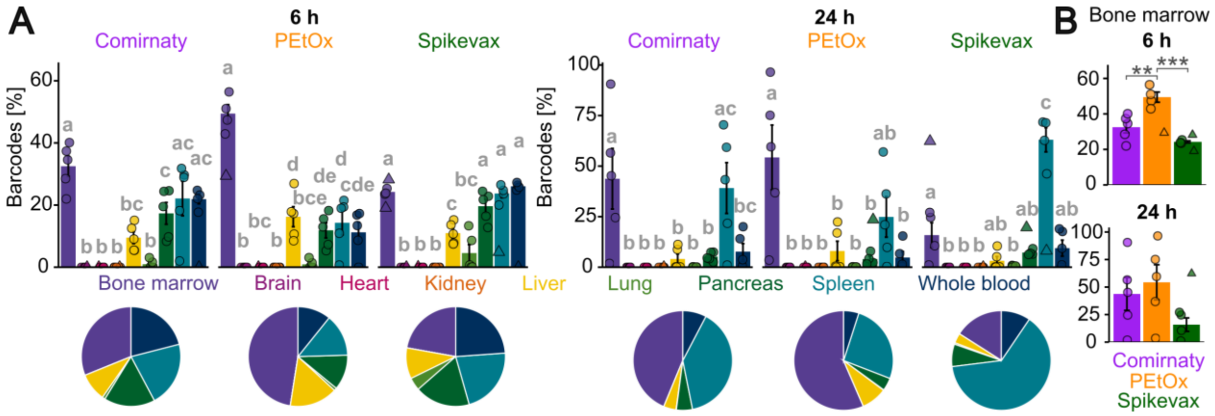

**Figure S9: LNP biodistribution results with filtering.** Plot analogous to **Figure 6** but with filtering points 1.5 × outside the interquartile range (from the first to the third quartile), prior to plotting and statistical analysis. Filtered points are shown as triangles (mean ± SEM; significant differences (p < 0.05) indicated with a (A) compact letter display (small letters a, b, and c) or (B) with stars, based on one-way ANOVA with post-hoc Tukey test: ** p < 0.01, *** p < 0.001).

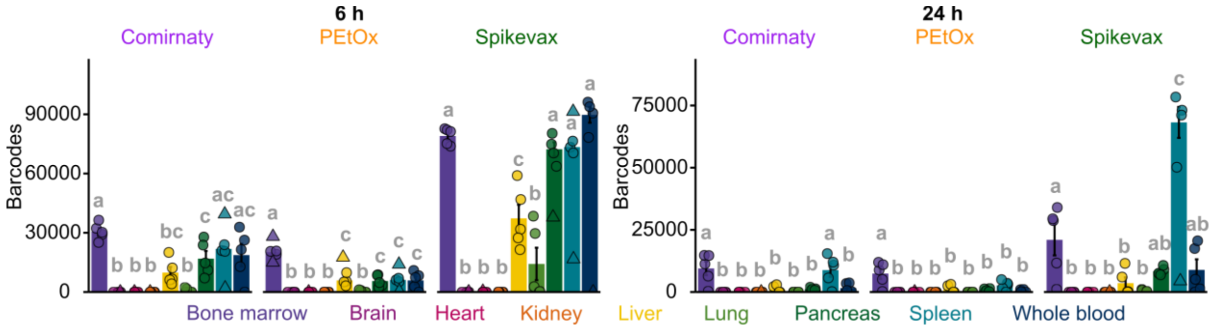

**Figure S10: Absolute DNA barcode amounts with filtering.** Plot analogous to **Figure S7** but with filtering points 1.5 × outside the interquartile range (from the first to the third quartile), prior to plotting (mean ± SEM; significant differences (p < 0.05) indicated with a compact letter display (small letters a, b, and c), based on one-way ANOVA with post-hoc Tukey test, performed separately for each formulation and time point). Filtered points are shown as triangles.

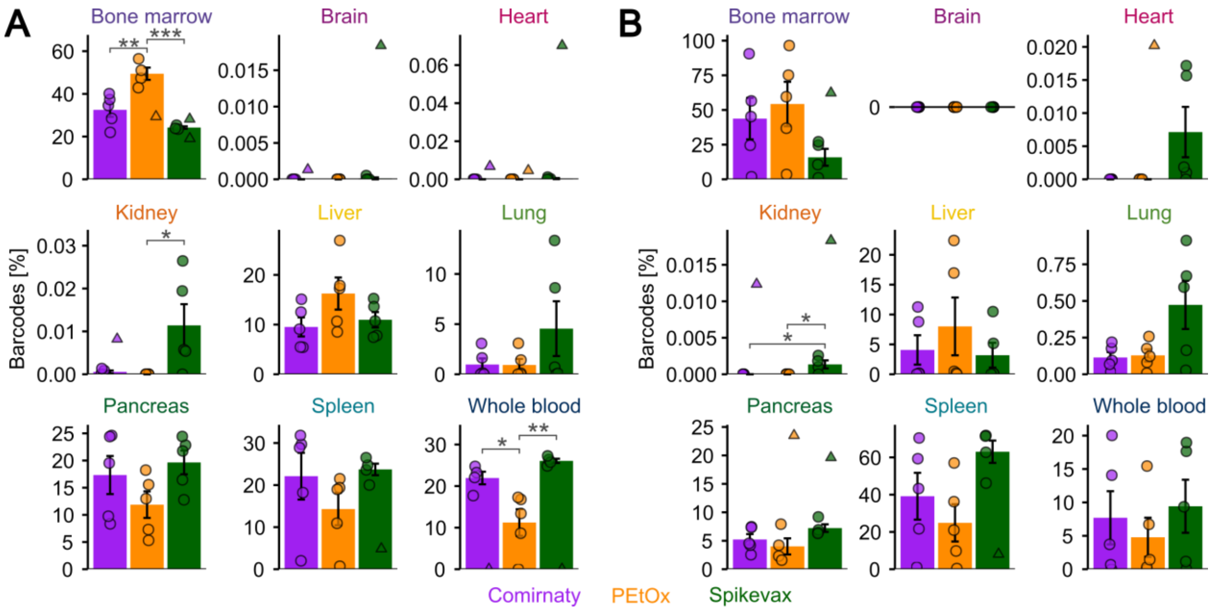


**Figure S11: DNA barcoding results per organ with filtering.** Plot analogous to **Figure S8** but with filtering points 1.5 × outside the interquartile range (from the first to the third quartile), prior to plotting and statistical analysis. Filtered points are shown as triangles (mean ± SEM; significant differences indicated with stars, based on one-way ANOVA with post-hoc Tukey test: * $p < 0.05$, ** $p < 0.01$, *** $p < 0.001$). (A) Results for 6 h. (B) Results for 24 h.

#### 7 References

[1] J. Leblond, N. Mignet, C. Largeau, M.-V. Spanedda, J. Seguin, D. Scherman, J. Herscovici, Lipopolythioureas: a new non-cationic system for gene transfer, Bioconjug. Chem., 18 (2007) 484–493.

[2] J. Schroeder, J. Ismail, C.T. Holick, J. Jungwirth, L. Klement, S. Hoeppener, C. Kosan, M. Schmidtke, B. Löffler, C. Weber, U.S. Schubert, C. Hoffmann, S. Schubert, C. Ehrhardt, Advancing influenza virus treatment: in vitro and ex vivo studies of PI3K inhibitor-loaded lipid nanoparticles, Mater. Today Bio, 35 (2025) 102587.

[3] D. Svergun, C. Barberato, M.H.J. Koch, CRYSOL – a program to evaluate X-ray solution scattering of biological macromolecules from atomic coordinates, J. Appl. Crystallogr., 28 (1995) 768–773.
